# MINFLUX nanoscopy delivers multicolor nanometer 3D-resolution in (living) cells

**DOI:** 10.1101/734251

**Authors:** Klaus C. Gwosch, Jasmin K. Pape, Francisco Balzarotti, Philipp Hoess, Jan Ellenberg, Jonas Ries, Stefan W. Hell

## Abstract

The ultimate goal of biological superresolution fluorescence microscopy is to provide three-dimensional resolution at the size scale of a fluorescent marker. Here, we show that, by localizing individual switchable fluorophores with a probing doughnut-shaped excitation beam, MINFLUX nanoscopy provides 1–3 nanometer resolution in fixed and living cells. This progress has been facilitated by approaching each fluorophore iteratively with the probing doughnut minimum, making the resolution essentially uniform and isotropic over scalable fields of view. MINFLUX imaging of nuclear pore complexes of a mammalian cell shows that this true nanometer scale resolution is obtained in three dimensions and in two color channels. Relying on fewer detected photons than popular camera-based localization, MINFLUX nanoscopy is poised to open a new chapter in the imaging of protein complexes and distributions in fixed and living cells.

---

While STED^1, 2^ and PALM/STORM^3, 4^ fluorescence microscopy (nanoscopy) can theoretically achieve a resolution at the size of a single fluorophore, in practice they are typically limited to about 20 nm. Owing to a synergistic combination of the specific strengths of these key superresolution concepts, the recently introduced MINFLUX nanoscopy^5^ can attain a spatial resolution of about the size of a molecule, conceptually without constraints from any wavelength or numerical aperture. In MINFLUX imaging, the fluorophores are switched individually like in PALM/STORM, whereas the localization is accomplished by using a movable excitation beam featuring an intensity minimum, such as a doughnut. The minimum ideally is a zero intensity point that is targetable like a probe^6^.

Concomitantly, it serves as a reference coordinate for the unknown position of the fluorophore in the sample, because the closer the minimum is to the fluorophore, the weaker is the emitted fluorescence per unit excitation power. If the excitation zero and the fluorophore coincided in space, no emission would occur. Yet the emitter’s position would be known in that case, since it must coincide with the well-controlled position of the zero. For the same reason, the smaller the mismatch between the two coordinates is, the fewer emitted photons are required to measure the emitter’s position. Hence, approaching a fluorophore with a position-probing excitation minimum shifts the burden of requiring many photons for localization from the feeble fluorescence to the inherently bright beam of molecular excitation, giving MINFLUX a fundamental edge over popular camera-based localization.

While the resolution and speed advantage of MINFLUX nanoscopy have been well documented^5, 7^, here we demonstrate that it also allows for three-dimensional (3D) imaging and simultaneous two-color registration, which are critically important for most life science applications. Moreover, we show that, in conjunction with photoactivatable fluorescent proteins, MINFLUX uniquely affords true nanometer resolution in living cells. The first realization of MINFLUX utilized static distances of the intensity zero to the fluorophore, which limited the imaged field to sub-100 nm extents^5^. In this work, we dynamically zoom-in on each fluorophore position, which not only renders the localizations essentially uniform and isotropic, but also facilitates the recording of extended areas and volumes.

## Results

### Basics on iterative fluorophore targeting with a local excitation minimum

The power of zooming-in on each fluorophore is readily illustrated for a fluorophore located at *x*_M_, within the interval *x*_0_ < *x*_M_ < *x*_1_ of size *L* = |*x*_1_ −*x*_0_|. When probing with an intensity zero that is flanked by a quadratic intensity profile, it is sufficient to measure the number of fluorescence photons *n*(*x*_0_) and *n*(*x*_1_) with the zero placed at the interval endpoints^5^. With *n*(*x*_0_) and *n*(*x*_1_) following Poissonian statistics, the minimum standard deviation of the localization, i.e. the Cramer-Rao bound (CRB) within the region *L*, is given by 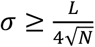, with *N* = *n*(*x*_0_) + *n*(*x*_1_). Contrasting the linear dependence on the interval size *L* with the inverse square-root dependence on the number of detected photons *N* shows that reducing *L*, i.e. zooming-in on the molecule, outperforms the wait for more photons.

This also becomes evident when approaching *x*_M_ in successive steps with stepwise reduced *L*_*k*_, such that *L*_*k*_ is chosen to match three times *σ*_*k*−1_, i.e. the uncertainty of the previous step. Thus, after *k* iterations we obtain a total of *N*_t_ = *k* ⋅ *N* detected photons and a CRB of

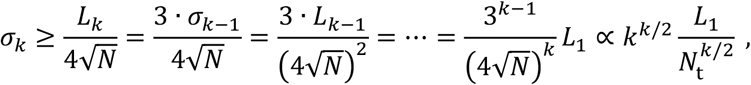

underscoring once more that it is more effective to increase the number *k* of iterations than that of the photons *N* per iteration. Already four steps (*k*=4) yield 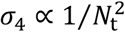, i.e. an inverse quadratic as opposed to an inverse square-root dependence on the total number of detected photons. More iterations yield an even higher order, accounting for the fact that the detected photons become more informative the closer the probing positions are to the molecule. In practice, this procedure is only limited by the background photons, which compromise the information on where to place the zero next.

### Realizing iterative MINFLUX and scalable fields of view

Our setup is a beam-scanning confocal fluorescence microscope featuring a visible excitation beam that can enter the objective lens with a flat wavefront, hence being regularly focused, as well as amplitude- and phase-modulated to form a doughnut in the focal region (Fig. 1a, Optical setup, Fig. S1). The excitation beam is co-aligned with a UV beam for activating individual emitters in a ~400 nm diameter region in the sample (Fig. 1a,e). Micrometer-range positioning or scanning of these co-aligned beams is performed with a piezo-based mirror, whereas fast microsecond nanometer-range targeting is carried out using an electro-optic beam deflector.

**Fig. 1.**
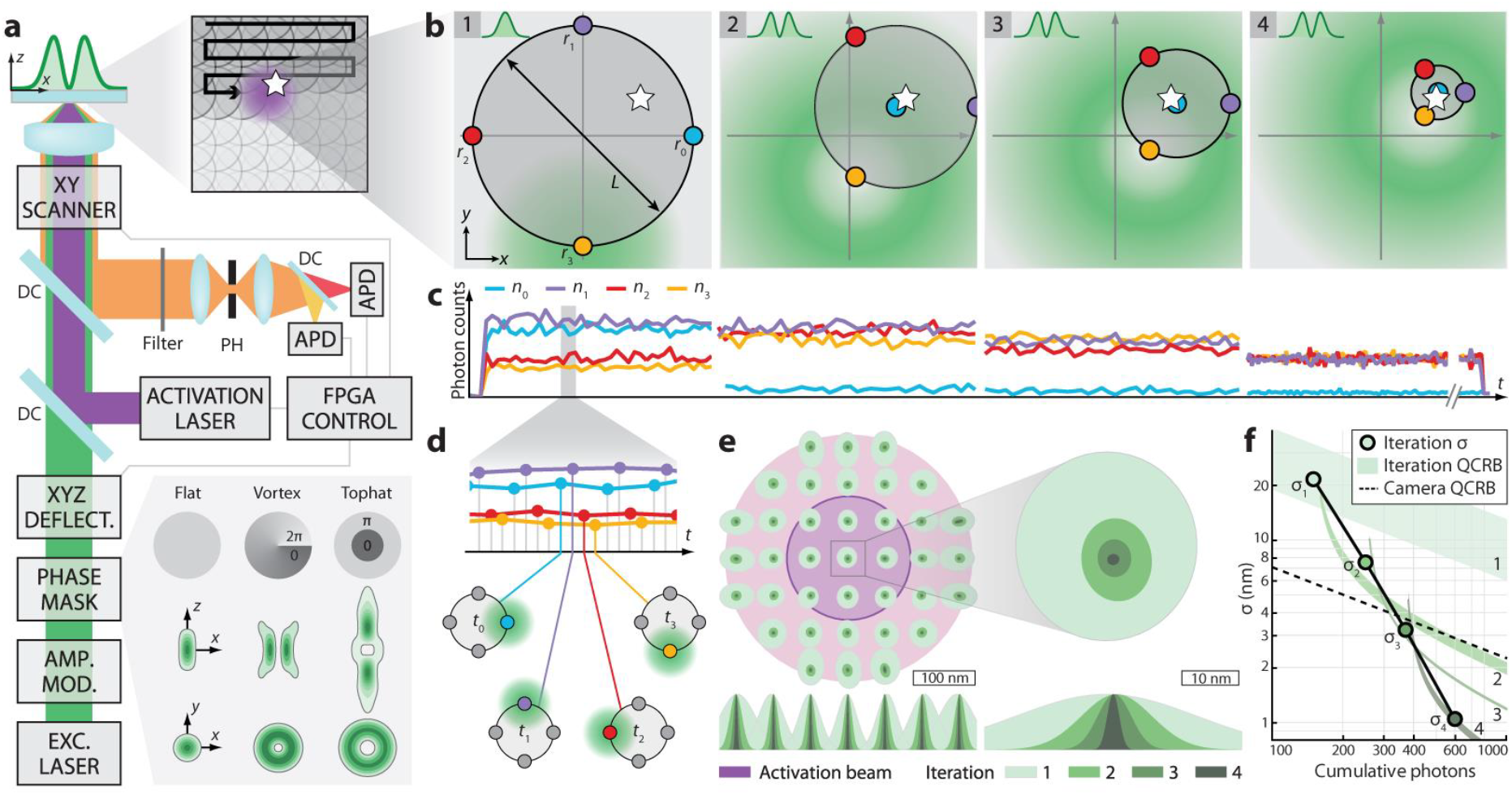
Iterative MINFLUX setup and localization. **a**, Setup. An excitation beam (green) is amplitude- and phase-modulated (bottom right inset: flat (regular focus), vortex (2D-doughnut), tophat (3D-doughnut)), electro-optically deflected in *x*,*y*,*z*, overlapped with a photo-activation beam (purple) and focused into the sample, after passing a piezo-actuated mirror for coarse *x*,*y*-scanning (top right inset, white star: activated molecule position). Fluorescence is de-scanned, deflected by a dichroic mirror (DC), filtered by a confocal pinhole (PH), split in two spectral ranges by another DC and detected by photon-counting avalanche photodiodes (APDs), all FPGA-board controlled. **b**, Iterative *xy*-localization by targeting the beam to four designated coordinates constituting the TCP (blue, purple, red, yellow and beam on yellow position in green). Step 1: regular focus, steps 2–4: 2D-doughnut. The TCP is re-centered and zoomed-in on the fluorophore (white star) in steps 2–4. **c**, Typical fluorescence counts for each iteration with the color indicating the targeted coordinate. **d**, Representation of the interleaved TCP measurement. **e**, Convergence of iterative localizations for molecules within the activation area (purple: 200 nm FWHM, 50% single molecule activation probability; pink: 2⋅FWHM, 95% activation probability). The covariance of each iteration (green shades) is represented as an ellipse of e^−1/2^ level. **f**, Progression of the spatially averaged localization precision *σ*_1_–*σ*_4_ for each iteration (green dots) with the corresponding CRBs (green shades) and the CRB for camera-based localization (dashed line). Conditions for (e– f): Photons: *N*_1_,=150 *N*_2_=100, *N*_3_=120 and *N*_4_=230; total *N*_t_=600. TCPs: *L*_1,flat_=300 nm, *L*_2,vortex_=150 nm, *L*_3,vortex_=90 nm, *L*_4,vortex_=40 nm.

Like all fluorescence nanoscopy concepts, MINFLUX nanoscopy relies on a fluorescence on-off transition for the separation of neighboring emitters^6^. We transiently activated a single fluorophore within the ~400 nm diameter activation region, localized it by MINFLUX and finally ensured that it went back to a lasting off-state. The same procedure was applied to the next fluorophore until a representative number of molecules was registered. For each localization, we targeted the zero to a set of coordinates around the anticipated fluorophore position, referred to as the targeted coordinate pattern (TCP). Forming a circle in the focal plane, the diameter *L* of the TCP is a measure of how well the TCP is zoomed-in on the molecule (Fig. 1b). In the first iteration, the beam was focused regularly, while in the succeeding steps it was modulated to form a 2D- or a 3D-doughnut, depending on whether we localized the fluorophore just in the focal plane (*x*,*y*) or in the sample volume (*x*,*y*,*z*). In each iteration, we calculated the newly anticipated fluorophore position based on collecting a defined number of photons in the fluorescence traces (Fig. 1c) produced at each of the targeted coordinates. The four-point TCP used for 2D-localization rendered the position estimator simple and unambiguous^5^. In the next iteration, the TCP was centered on the new position and the diameter *L* decreased according to the new precision estimate, thus bringing the zeros closer to the molecule. To compensate for the associated reduction in excitation intensity and to maintain the fluorescence flux, we increased the excitation power in each step. The smallest *L* in the last iteration step, meaning how well we could zoom-in on the fluorophore in practice, was determined by the concomitantly decreasing signal-to-background ratio.

We first simulated the procedure by randomly generating multinomial photon counts for different molecule positions, applying four iteration steps, and zero background (see Simulations and optimization of iterative strategy). On average, our iteration procedure (Fig. 1b) achieved a localization precision of *σ*_1D_~1 nm with only *N*_t_=600 photons (Fig. 1e). In contrast to static (non-iterative) MINFLUX localization^5^, the precision obtained here was largely isotropic and independent of the emitter position within the activation area (Fig. 1e). In our simulation, we selected different photon numbers in each step according to the required precision, with the last step featuring the largest photon number. The precision still largely followed the anticipated 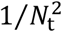 behavior. Starting with a lower photon efficiency, our iterative MINFLUX scheme surpassed the quantum CRB (QCRB)^8^ of lateral precision for camera-based localization with *N*_t_~330 photons (Fig. 1f).

For imaging, we defined a state machine on a field-programmable gate array (FPGA) board that recognized single activated fluorophores and localized them iteratively (see Experiment control software). Individual emitters were identified by segmenting their emission trace and their position was obtained by maximum-likelihood estimation (see Data analysis). First, we imaged Nup96, a protein of the nuclear pore complex (NPC), labelled with the organic fluorophore Alexa Fluor 647 using SNAP-tag in fixed mammalian (U-2 OS) cells^9^ (Fig. 2a, see U-2 OS NUP96-SNAP/mMaple). Images of several micrometers in size were acquired using MINFLUX with five iteration steps (*N*_1_=100, *N*_2_=100, *N*_3_=150, *N*_4_=300, *N*_5_~2000 and *L*_1,flat_=300 nm, *L*_2,flat_=300 nm, *L*_3,vortex_=150 nm, *L*_4,vortex_=100 nm, *L*_5,vortex_=50 nm). Based on these parameters and the background, we expected a localization precision below 1 nm in standard deviation. The overall fluorescence rate of typically ~50 kHz allowed a complete localization within ~40 ms. To ensure a single active fluorophore per activation area, 0.5–5 µW of UV light was applied in pulses of 5 ms until a fluorescent molecule appeared. After activation and registration of a single Alexa Fluor 647 molecule, the iterative scheme took around 10 ms to reach the final iteration. The photon traces and the TCP re-adjustment were constantly monitored during recording. Altogether, the acquisition of a single NPC required about 1– 2 minutes. Our experiments demonstrated that iterative MINFLUX clearly resolved the eightfold symmetry of Nup96 in single NPCs (Fig. 2a), distributed along a ring of 107 nm in diameter^9, 10^. In fact, the localizations typically formed eight clusters of roughly 2–4 localization subclusters, displayed as the sum of Gaussian distributions, one for each localization (see Image rendering), revealing, most likely, individual Nup96 proteins through their individual fluorescent markers. This molecular scale resolution is obtained in various images of the nuclear envelope, demonstrating that iterative MINFLUX nanoscopy can accommodate fields of any size and shape, just like any other scanning confocal microscope.

**Fig. 2.**
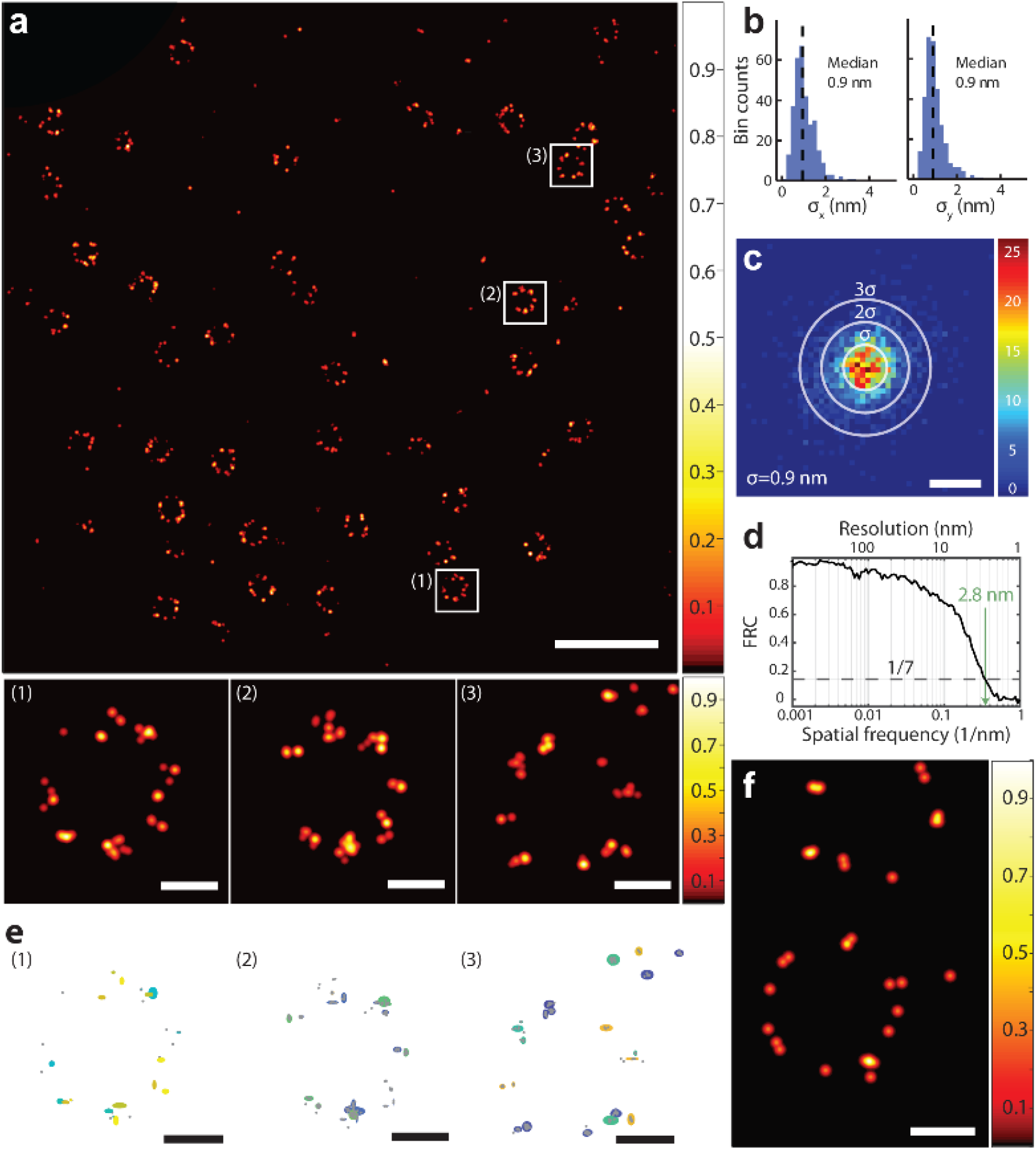
MINFLUX nanoscopy in fixed and living cells. **a**, MINFLUX nanoscopy of a U-2 OS cell expressing Nup96-SNAP labelled with Alexa Fluor 647. Single molecule fluorescence events were split into bunches of 2000 photons, yielding localizations that are filtered and displayed according to the methods sections Event filtering, Image rendering and Fig. S2. Zoomed-in excerpts (below) show single nuclear pores, with each dot highlighting groups of localizations representing individual proteins through their fluorescent labels. **b**, Histograms of standard deviations of localizations from single molecules in *x* and *y*. Only molecules providing >4 localizations per trace were considered. **c**, Histogram of the distance of a localization to the mean position of a single emitter. The ellipses are displayed with major axes of 2*σ*, 4*σ* and 6*σ* length respectively, with *σ* being determined in a 1D Gaussian fit. **d**, Fourier ring correlation curve for the image in (a). **e**, Renditions of insets of (a) with 3*σ* ellipses for each molecule. **f**, MINFLUX image of nuclear pores in living U-2 OS cell expressing NUP96-mMaple; average of ~1600 detected photons per localization. Scale bars: 500 nm (a), 50 nm (a (inlets), e, f), 2 nm (c).

To provide quantitative resolution measures, we applied three different criteria (Fig. 2b–d, see Performance metrics). First, we calculated the standard deviation *σ* for four and more localizations per fluorophore, where each localization utilized ~2000 photons of the last iteration, extracted from over 300 single fluorophore events. The resulting distribution of *σ* featured a median of ~1 nm (Fig. 2b). Another resolution assessment was gained by subtracting the mean localized position of an emission train from all of its single localizations (Fig. 2c). Assuming a Gaussian distribution, we obtained a precision error *σ*_1D_=0.9 nm. In the third approach, we calculated the Fourier ring correlation (FRC) curve^11, 12^ of the localization data (Fig. 2d), which better captures the influence of setup instabilities on the effective resolution. The threshold value of 1/7 gave Δ_FRC_~3 nm as a resolution estimate. Simulating localizations at the expected Nup96 positions with a fixed localization standard deviation revealed that this experimental FRC resolution Δ_FRC_ corresponds to a localization precision of roughly 1 nm. To represent the obtained precision in an actual image, we displayed individual localizations with covariance ellipses of 3*σ* for each emission (Fig. 2e).

Since Alexa Fluor 647 imposes a number of restrictions on the sample preparation, most notably incompatibility with living cells, we next demonstrated that MINFLUX nanoscopy is viable with photoactivatable fluorescent proteins. We resolved the eightfold symmetry of Nup96 tagged with the photo-convertible protein mMaple (see U-2 OS Nup96-SNAP/mMaple and Tab. S1) in both fixed and living cells (Fig. 2f). In particular, the live-cell recording proves that imaging of protein complexes in the interior of living cells is possible with nanometer resolution.

### 3D MINFLUX imaging with isotropic resolution

Dissection of macromolecular complexes calls for 3D-resolution and hence for usage of the 3D-doughnut (Fig. 3a). In our 3D localization algorithm, we probe the molecule position with three different TCPs. First, the *xy*-position is probed with a regularly focused beam, so that the central *z*-axis of the 3D-doughnut is positioned as closely as possible to the fluorophore. Next, the 3D-doughnut is targeted in two positions above and below the anticipated fluorophore position. Last, three coordinate pairs on the *x*-, *y*- and *z*-axis and also the center coordinate of this arrangement are addressed, resulting in a 3D position estimate. A simulation of this 3D-MINFLUX algorithm (Fig. 3b) showed that 1000 detected photons were sufficient to cover a ~400 nm diameter volume with a largely homogenous and isotropic 3D-precision of ~1 nm (Fig. 3c-d). Again, the increase in precision with the total number of detected photons proved to be steeper than 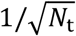 (Fig. 3e). Comparing. the result with the QCRB for the precision of camera-based localization shows that 3D-MINFLUX can outperform all camera-based approaches, including multiple-objective lens arrangements.

**Fig. 3.**
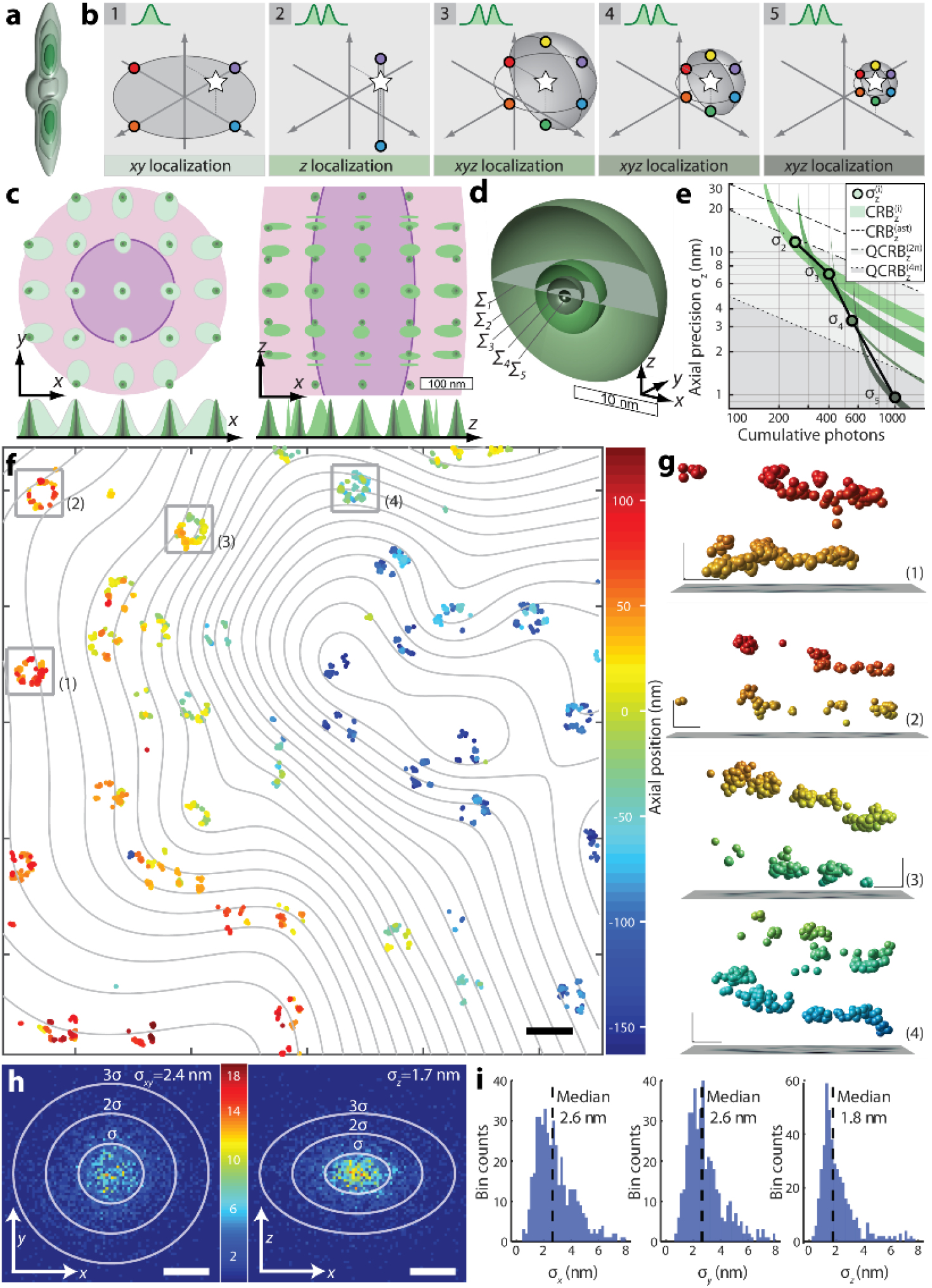
Iterative 3D MINFLUX localization yields isotropic nanometric precision. **a**, 3D-doughnut with selected isointensity surfaces. **b**, Iterations. Step 1: *xy*-localization with regularly focused beam (intensity profile top left). Step 2: *z*-localization with 3D-doughnut. Steps 3–5: *x*-,*y*-, and *z*-localization with 3D-doughnut, shrinking the TCP each time. **c**, Simulation of localization convergence in *xy* (left) and *xz* (right) for molecules within the activation region (purple: 200 nm FWHM_*xy*_ and 600 nm FWHM_*z*_ with 29% activation probability, pink: 2 FWHM with 86% activation probability). The covariance of each iteration (green shades) is represented as an ellipse at level e^−1⁄2^. **d**, 3D representation of the step-wise localization precision in the central region in c. **e**, Progression of average *z*-precision in c (dots), showing the Cramér-Rao bound (CRB) for camera-based astigmatic localization, the Quantum CRB (QCRB) for a camera localization in a 2*π* and 4*π*-arrangement (dashed lines) along with MINFLUX CRB in each iteration (green shades). Steps for c-e: Photons *N*_1_=150, *N*_2_=100, *N*_3_=150, *N*_4_=150, and *N*_5_=450; total *N*_t_=1000. Step 1: regular focus with *L*_1,flat_=300 nm. Steps 3–5: 3D-doughnut with *L*_2,tophat_=400 nm, *L*_3,tophat_=150 nm, *L*_4,tophat_=90 nm, *L*_5,tophat_=40 nm. **f**, MINFLUX image of nuclear pore complexes of a fixed cell (Nup96-SNAP-Alexa Fluor 647); *z*-coordinate is color-coded. Contour lines indicate the nuclear envelope. **g**, *z*-sections of single nuclear pores in (f). **h**, Localization precision for 3D imaging in *xy* (left) and *xz* (right). **i**, Histograms of *x*, *y*, *z* standard deviation. Scale bars 200 nm (f), 20 nm (g), 5 nm (h).

In our setup, the axial position of the 3D-doughnut is controlled with an electrically tunable lens, allowing to refocus within 50 µs. Imaging Nup96-SNAP-Alexa Fluor 647 (Fig. 3f, Fig. S3, Sample preparation, Video S1), 3D-MINFLUX now discerns the cyto- and nucleoplasmic layers of Nup96 in single pore complexes (Fig. 3g). As the cell lies flat on a cover slip, these layers are typically parallel to the focal plane and ~50 nm apart in the z-direction^9^. 3D-MINFLUX also recovered the curvature of the nuclear envelope extending ~300 nm in depth within the acquired region (Fig. 3f). The localizations featured one-dimensional (1D) standard deviations *σ*_*xy*_=2.4 nm and *σ*_*z*_=1.7 nm (Fig. 3h). Localizations from single events exhibited median standard deviations *σ*_*xy*_=2.6 nm and *σ*_*z*_=1.8 nm respectively (Fig. 3i). Note that the refractive index mismatch at the glass-water interface causes a slight difference between the *xy* and *z* localization precision, since *L*_*z*_ of the TCP is slightly compressed due to this mismatch^13^, increasing the MINFLUX localization precision accordingly (see Optical setup).

Next, we imaged the protein PSD-95 in dissected hippocampal neuron cultures from transgenic mice expressing a fusion protein of the Halo-Tag enzyme connected to the protein C-terminus^14^ (Fig. 4a–b, Fig. S4, Hippocampal cultured neurons (PSD-95) and Video S2). PSD-95 putatively plays a key role in anchoring and re-arranging glutamate receptors in the post-synaptic membrane^15, 16^. 3D-MINFLUX nanoscopy suggests that PSD-95 is arranged along a slightly curved surface of 100–400 nm side length. The spots of a high localization density exhibit a standard deviation of ~4–6 nm, with the localization precision being estimated to *σ*_1D_~2–3 nm (Fig. 4c). The average distance between nearest-neighbor clusters is ~40 nm. While superresolution techniques were repeatedly used to elucidate the organization of post-synaptic proteins^17^, virtually all superresolution studies were carried out in 2D. Owing to its isotropic resolution, 3D MINFLUX now opens up entirely new possibilities for studying synaptic protein organization.

**Fig. 4.**
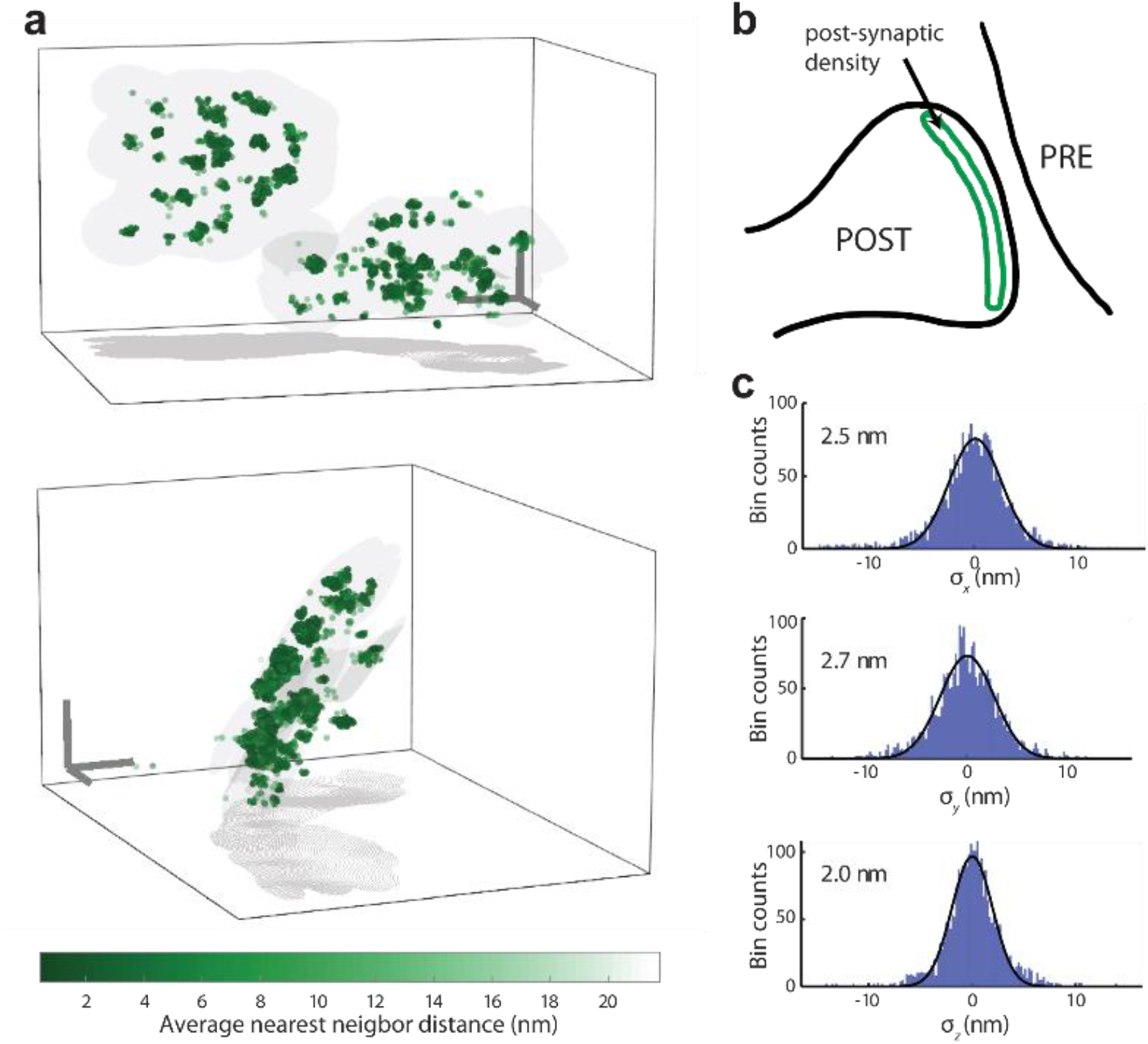
MINFLUX imaging of the post-synaptic protein PSD-95 with 3D-resolution of ~2–3 nm. Sample: hippocampal neurons from transgenic mice expressing PSD-95-Halo conjugated to Alexa Fluor 647 after fixation. **a**, 3D-MINFLUX image with color indicating the 3D density of molecular localizations. PSD-95 appears in clusters distributed on a curved surface. **b**, Sketch of PSD-95 occurrence at the postsynaptic site. **c**, Histograms of distances of localizations to the mean molecule position in *x*, *y* and *z*, revealing an isotropic 3D localization precision of the individual fluorophores of 2.0–2.7 nm. Scale bar: 100 nm (a).

### Multicolor MINFLUX imaging

Fluorophore species sharing the same excitation wavelength but having measurable differences in their emission spectra, such as Alexa Fluor 647, CF660C, and CF680^18^ can be separated by spectral classification. Hence, we split the fluorescence at 685 nm into two spectral fractions using a dichroic mirror, and detected both fractions with photon counting detectors. While localization was performed adding up the photons from both detectors, comparing the counts of each spectral fraction enabled fluorophore classification (see Multicolor classification). The classification was refined using a principal component analysis on the spectral fractions from all MINFLUX iterations and selecting a classification threshold based on the distribution of the first principal component. We first tested two-color 2D MINFLUX imaging on a DNA origami labelled with Alexa Fluor 647 and CF660C molecules, arranged at distances of ~12 nm (Fig. 5a, Fig. S4, see DNA origami). MINFLUX nanoscopy correctly recovered the labelling sites (Fig. 5b,d) and distinguished the two fluorophore species without overlap (Fig. 5c). Moreover, chromatic distortions between the two color channels are excluded on principle grounds, since the localization is performed by one and the same excitation beam. This is in contrast to camera-based localization, where nanometer multicolor co-localization is affected by chromatic aberrations of the optical system.

**Fig. 5.**
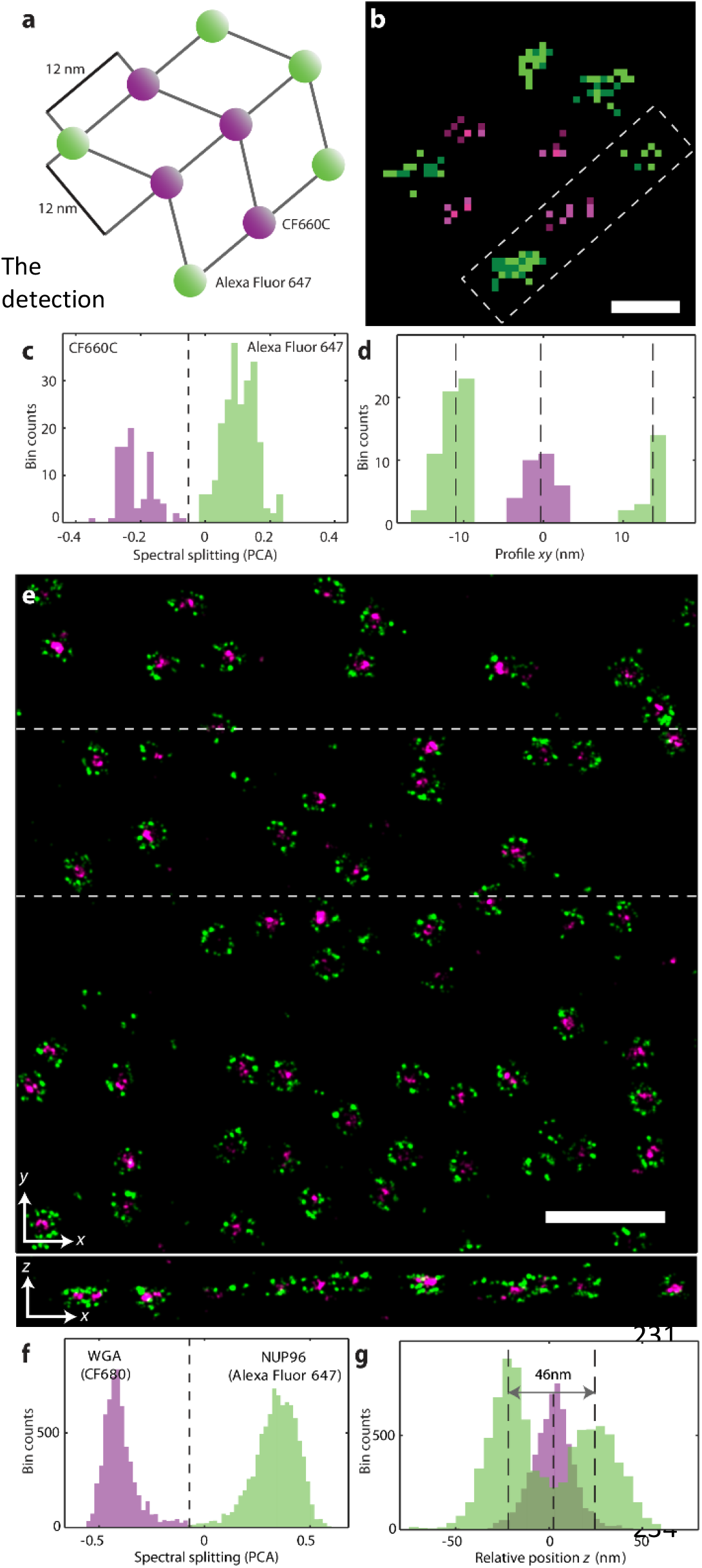
Two-color MINFLUX nanoscopy in 2D and 3D. **a**, Scheme of a DNA origami labelled with Alexa Fluor 647 and CF660C with ~12 nm intermolecular distances. **b**, Histogram of the localizations acquired with 2D MINFLUX. Alexa Fluor 647 is shown in green, CF660C in magenta. **c**, Spectral separation of the two fluorophore species. abscissa is the principal component of the probability on the spectral channel with *λ*<685 nm for all iterations. The spectral separation (dashed line) minimizes overlap between the two species. **d**, Histogram of localizations along the box in b. The positions are well resolved with a distance of the outer fluorophores of ~25 nm. **e**, Two-color image of U-2 OS cell expressing NUP96-SNAP. The SNAP-tag was labelled with Alexa Fluor 647. Additionally, wheat germ agglutinin (WGA) conjugated to CF680 stained the center of the NPC. **f**, Spectral distribution of the two fluorophore species with choice of color separation (dashed line). **g**, z-distribution of the two fluorophore species. After considering the nuclear envelope shape (see Fig. 3f) for each nuclear pore position, a distance of 46 nm is found between the cytoplasmic and nucleoplasmic layers of Nup96; WGA is aggregated at the pore center. Scale bars 10 nm (b) and 500 nm (e).

Finally, we acquired two-color 3D-images of the nuclear pores in the U-2 OS cell by labeling the nuclear pore center with wheat germ agglutinin (WGA) conjugated to CF680 in addition to the Nup96-SNAP Alexa Fluor 647 decoration^19^. We indeed observed the CF680 residing inside the Nup96 octamer both laterally and axially (Fig. 5e,g, Video S3). Although the emitters featured a broader spectral distribution in the cell than in the origami, we could classify the labels with high fidelity (Fig. 5f). We also quantified the distribution of both emitter species along the z-axis by taking into account the nuclear envelope 3D curvature, which was found through a spline interpolation of the pore positions (see Nuclear pore complexes). Thus, we recovered a distance of ~46 nm between the Nup96 layers with the WGA distribution centered in between (Fig. 5g), underscoring once more the 3D capability of MINFLUX nanoscopy.

## Discussion and Conclusion

We have shown that MINFLUX nanoscopy provides nanometer resolution (1–3 nm in standard deviation) in three dimensions, on arbitrary fields of views, in living cells, and using multiple color channels. Yet, the full potential of this method has not been reached. Ongoing and future developments will cut down the current recording time of tens of minutes per 500 localizations. This can be achieved by optimizing the activation procedure, registering more than just one molecule per localization, and minimizing the fluorophore-to-zero average distance. The latter should be possible through background reduction by time-gated detection and multiphoton excitation. Note that the excitation powers used in this work are on the order of 20–60 µW, so that the intensities are comparable to those in biological confocal microscopy, where the power is confined to an area three times smaller than that of a doughnut beam.

Since MINFLUX utilizes a scanning read-out, the recording time of MINFLUX nanoscopy inevitably scales with the imaged area or volume. However, an advantage of scanning is that one can adapt the activation rate to the local fluorophore concentration to save time. Another avenue is to parallelize the scanning process using sets of doughnuts or standing waves. Furthermore, one can utilize different on-off switching processes, such as the transient binding of fluorescent molecules like in PAINT^20^ or DNA-PAINT^21^. Note that for a PAINT-like sample preparation, the speed and precision advantages provided by MINFLUX persist.

With lens-based fluorescence microscopy having finally reached true nanometer resolution, it is important for life scientists to bear in mind that fluorescence microscopes map nothing but the fluorophores; once they have fulfilled this task, they have accomplished their mission. Hence, any conclusion going beyond that, especially at sub-10 nm length scales, has to take the size and orientation of the fluorophore label into account, making the labeling method ever more crucial. MINFLUX offers more flexibility in this regard because, by requiring fewer detected photons than camera-based localization, a larger range of fluorophores and labeling procedures should be viable. For this fundamental reason, as well as its 3D imaging performance, MINFLUX nanoscopy is bound to be a cornerstone, if not the vanguard, of nanometer scale fluorescence microscopy in the years to come.

## Acknowledgements

We thank the following MPI colleagues: Elisa D’Este for support with biological sample optimization; Mark Bates for discussions about single molecule techniques; Haisen Ta for help with DNA origami; Dirk Kamin and Ina Herfort for support with cultured neurons. We acknowledge Seth Grant (University of Edinburgh) for providing the PSD-95-Halo mouse model. Jervis Thevathasan, Bianca Nijmeijer, Moritz Kueblbeck and Ulf Matti (all EMBL) made the Nup96 cell lines. P.H. and J.R. acknowledge the European Research Council (ERC CoG-724489) and the Human Frontier Science Program (RGY0065/2017, J.R.).

## Author Contributions

K.C.G., J.K.P. and F.B. designed and built hardware, programmed software, performed experiments and data analysis with input from S.W.H. K.C.G, J.K.P and P.H. prepared samples. P.H. and J.R. proposed the Nup96 cell line as resolution test sample. J.E. made the Nup96 cell line. F.B. carried out theoretical analysis of MINFLUX with K.C.G and co-supervised the project. S.W.H. supervised the project and is responsible for its conceptual thrust. K.C.G., J.K.P., F.B. and S.W.H. wrote the manuscript.

## Materials and methods

### Experimental setup

#### Optical setup

We used an optical setup (Fig. S1) based on a previously described microscope^1, 2^, having three different illumination modalities provided through separate optical paths: (i) widefield excitation (488 nm, 560 nm, 642 nm), (ii) regularly focused excitation (560 nm or 642 nm) or focused activation (405 nm) and (iii) phase-modulated excitation (560 nm or 642 nm) leading to a 2D- or 3D-doughnut in the focal region. For all excitation beams, we employed an acousto-optic tunable filter (AOTF) for slow power modulation and wavelength selection. Switching between the regularly focused and 2D-/3D-doughnut excitation was performed with electro-optical modulators (EOM). To laterally position the MINFLUX beams within the TCP and to approach the molecules iteratively, we employed electro-optical deflectors (EOD); they were differentially driven with two pairs of identical amplifiers. To image larger fields of views, we scanned the focused activation beam and the TCP across the sample using a piezo-based tip-tilt mirror. An electrooptically actuated varifocal lens (VFL) positioned the 3D-doughnut beam axially. The sample was translated using a manual as well as a piezo-driven sample stage. 2D- and 3D-doughnut generation was accomplished with a spatial light modulator (SLM). Applying an achromatic λ/4 retarder plate for circular beam polarization helped minimizing the intensity at the doughnut minimum. To ensure a deep minimum, we measured the aberrated wavefront using a pupil segmentation based scheme^3^. A 1.4 numerical aperture oil immersion lens focused the excitation light into the sample and collected the fluorescence light. The tip-tilt mirror de-scanned the emitted light, which subsequently passed through a quad-band dichroic mirror. Electrically driven flip-mirrors allowed the selection between (i) a detection path for acquiring a fluorescence overview image of the sample with an EMCCD camera and selecting a region of interest; (ii) a large-area detector for measuring the point spread function (PSF) of the excitation beam; (iii) a confocal pinhole (multi-mode-fiber) of 500 nm (sample units) for MINFLUX fluorescence acquisitions.

An active stabilization system ensured nanometer stability of the sample. For lateral stabilization, we imaged micrometer-sized scattering gold nanorods onto a camera using a spectrally filtered white-light laser source (~950–1000 nm). An infrared laser beam (905 nm) illuminated the sample in total internal reflection mode, so that the axial position was accessible using another camera. A PID controller commanded the piezo-stage to compensate the movements. Since MINFLUX relies on the precise knowledge of the shape and position of the excitation beam, we mapped the applied scanner voltage to a physical position in the sample by calibrating the *xy*-scanners for both excitation lines^1^. Additionally, we calibrated the axial beam displacements (induced by the VFL) by measuring the axial position of the excitation PSF for different input voltages. We also took the refractive index mismatch between the coverglass and the sample into account. To this end, we performed a 3D-MINFLUX measurement to determine the position of two Alexa Fluor 647 molecules on a DNA nanoruler (GATTA-STED 3D 90R – custom, GATTAquant GmbH, Hiltpoltstein, Germany). We scaled the estimated *z* position to match the expected inter-molecule distance. We confirmed the scaling factor of 0.7 in simulations^4^ and applied it in post-processing to correct all estimated *z* localizations. Fluorescent microspheres (see Fluorescent microsphere sample) were used to re-examine the alignment of the microscope on a daily basis. To this end, we verified the coalignment of the activation beam, the regularly focused and the doughnut-shaped excitation beam. Moreover, we adjusted the confocal detection to coincide with the excitation volume. We measured the PSF of the excitation beams by scanning an area or volume around the doughnut minimum with the EOD or the tip-tilt mirror (*xy*) and the sample stage (*z*) respectively.

#### Experiment control software

Similarly to our previously published works^1, 2^ we used custom programs written in LabView (National Instruments, Austin, TX, USA) for control of the experimental setup and data acquisition. We implemented the control of hardware components using data acquisition boards (NI PCIe-6353), an FPGA board (NI USB-7856R), and through direct communication with devices (e.g. USB or serial port). The software package consisted of four types of programs: iterative MINFLUX FPGA core, iterative MINFLUX PC host, system stabilization, and auxiliary device control. The iterative MINFLUX FPGA core served the following purposes: control of beam deflectors and amplitude modulators; selection of beam shape and type (activation/excitation); photon registration via avalanche photo diodes (APDs); online data processing; and position scanning for image acquisition. This program performed all tasks and processes described in Data acquisition. The MINFLUX FPGA core and PC host communicated via USB, delivering control commands from the host PC to the FPGA and streaming the acquired data and state information from the FPGA to the host PC.

### Live position estimators

#### Position estimation for doughnut exposures

For 1D, 2D and 3D position estimation with a doughnut-shaped excitation beam, we used a modified version of the previously developed modified least mean square estimator (mLMSE)^1^. For position estimation in three dimensions, we performed a natural extension of the estimator resulting in the same expression as eq. (S51) in our reference^1^ but with the beam positions 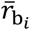 now being three-dimensional vectors:

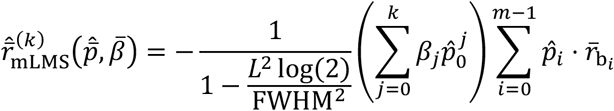

where *m* is the number of exposures (targeted coordinate positions), *k* is the estimator order, {*β*_*j*_}are the estimator parameters, *L* is the beam separation, FWHM is the full width at half maximum of a regularly focused beam (FWHM =360 nm in this work), and 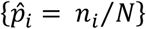 are the measured relative photons counts for the different exposures. We used an estimator order of *k* = 1. The estimator parameters *β*_*j*_ were optimized numerically using a typical measured shape of the 2D doughnut or an ideal quadratic function approximating the minimum of the 3D doughnut and assuming a typical experimental signal-to-background ratio. For optimization of {*β*_*j*_} we used the Matlab function *fmincon* to minimize the mean bias of the mLMSE position estimate in a circular region (for 2D) or spherical volume (for 3D) with a diameter of 0.8 *L*, with the beam separation *L*. We used the obtained position estimator parameters for the live position estimation in the MINFLUX FPGA core as well as for optimization of the iterative MINFLUX strategy.

#### Position estimation for regularly focused exposures

Position estimation using a regularly shaped excitation beam was performed using an estimator derived from the previously published estimator for position estimation in 1D using a TCP with a regularly focused beam and two exposures (see eq. (S40) in ^1^). We modified the molecule position estimator 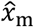 to take into account background signal. The estimator used in this work takes the form

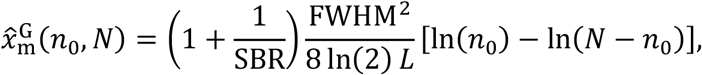

where *n*_0_ is the number of photons in the first exposure, *N* is the total number of detected photons, *L* is the beam separation of the two probing positions, FWHM is the full width at half maximum of the excitation beam (FWHM=300 nm in this study) and SBR is the signal-to-background ratio. We used this estimator for live position estimation in the MINFLUX FPGA core as well as for optimization of the iterative MINFLUX strategy. The SBR was kept fixed for each measurement assuming a typical value for the sample of study.

### Simulations and optimization of iterative strategy

Different iterative MINFLUX schemes were investigated and optimized through numerical simulations using Matlab (MathWorks, Natick, MA, USA). We simulated the complete iterative localization scheme for a set of molecules within the volume covered by the activation beam (360×360×360 nm), with the goal of finding the optimal number of iterations *K*, size of the TCP *L*_*k*_ and number of photons to use *N*_*k*_. The simulation replicated the live estimations performed by the FPGA hardware, while the final localization was performed via the MLE, which is used in post-processing. By minimizing the average localization error for all simulated molecule positions, the optimal iterative parameters {*K*, *L*_*k*_, *N*_*k*_} were found. The simulation used the following beam shapes:

| Type | Function | Variables |
| --- | --- | --- |
| Gaussian detection | $f(\vec{x}) = \frac{e^{\left(-\frac{(x-x_0)^2}{2\sigma_{\text{DetXY}}^2}\right)} \cdot e^{\left(-\frac{(y-y_0)^2}{2\sigma_{\text{DetXY}}^2}\right)} \cdot e^{\left(-\frac{(z-z_0)^2}{2\sigma_{\text{DetZ}}^2}\right)}}{(2\pi)^{3/2} \cdot \sigma_{\text{DetXY}}^2 \cdot \sigma_{\text{DetZ}}}$ | $\sigma_{\text{DetXY}}: \frac{500 \text{ nm}}{2\sqrt{2\log(2)}}$<br>$\sigma_{\text{DetZ}}: \frac{1000 \text{ nm}}{2\sqrt{2\log(2)}}$ |
| Focused excitation | $f(\vec{x}) = \frac{e^{\left(-\frac{(x-x_b)^2}{2\sigma_{\text{GaussXY}}^2}\right)} \cdot e^{\left(-\frac{(y-y_b)^2}{2\sigma_{\text{GaussXY}}^2}\right)} \cdot e^{\left(-\frac{(z-z_b)^2}{2\sigma_{\text{GaussZ}}^2}\right)}}{(2\pi)^{3/2} \cdot \sigma_{\text{GaussXY}}^2 \cdot \sigma_{\text{GaussZ}}}$ | $\sigma_{\text{GaussXY}}: \frac{330 \text{ nm}}{2\sqrt{2\log(2)}}$<br>$\sigma_{\text{GaussZ}}: \frac{800 \text{ nm}}{2\sqrt{2\log(2)}}$ |
| Doughnut 2D | $f(\vec{x}) = f_{\text{PSF2d}}(x - x_b, y - y_b) \cdot \frac{e^{\left(-\frac{(z-z_b)^2}{2\sigma_z^2}\right)}}{\sqrt{2\pi} \cdot \sigma_z}$ | $\sigma_z: \frac{1000 \text{ nm}}{2\sqrt{2\log(2)}}$<br>$f_{\text{PSF2d}}: \text{experimental 2D PSF}$ |
| Doughnut 3D | $f(\vec{x}) = (x - x_b)^2 + (y - y_b)^2 + (z - z_b)^2$ | |

### Data acquisition

Before starting a MINFLUX measurement, we selected a region of interest and moved the sample to the imaging plane based on a widefield fluorescence acquisition. We defined scanning points in the selected region, located either on a rectangular grid or in a manually defined arrangement. Using a focused excitation beam with high intensity (100–200 µW entering the back focal plane of the objective lens), we transferred the fluorescent molecule population into a long-lived non-fluorescent state. We conditionally applied light of 405 nm wavelength (0.5–5 µW) to photo-activate single molecules. Subsequently, we probed the presence of an emitting fluorophore by illuminating with excitation light using the configuration defined for the first iteration (typically 0.1 ms minimal time until collecting 40–100 photons) and comparing the low-pass filtered photon count rate (filter constant 1–10 ms) to a pre-defined threshold (5–50 kHz). Upon detection of a fluorescent molecule, we did not return to the activation step and continued the iterative MINFLUX acquisition scheme.

During the iterations, we applied excitation powers in the range of 20–60 µW in the backfocal plane comparable to confocal microscopy. The MINFLUX scheme ended (i) upon the photon count rate falling below a threshold pre-defined for each iteration, meaning the molecule turned off (ii) upon reaching a certain photon number in the last iteration. In case (i), activation light was applied. In case (ii), we restarted the MINFLUX iterations. After a pre-defined time with no single molecule events (0.3–20 s) or a certain amount of activation pulses (10000–30000 pulses corresponding to 50–150 ms of 405 nm light illumination), the tip-tilt mirror moved the illumination beams to the next scan position. We selected the scan positions in a rectangular pattern with distances of 200–250 nm, ensuring a homogenous activation of the whole scan area. As a result, regions covered in neighboring scan positions were overlapping, so that a molecule between the scan positions could be approached starting from any of the neighboring positions.

### Data analysis

#### PSF evaluation

We measured the shape of the excitation point spread function (PSF) with samples of immobilized fluorescent microspheres (FluoSheres®, 0.02 µm, dark red fluorescent; Thermo Fischer Scientific Inc., Waltham, MA, USA). For measurements in two dimensions, data acquisition and processing of the PSF followed a previously published protocol^1^. For measurement in three dimensions, we assumed the PSF to take the form *I*(*x*, *y*, *z*) = *a* ⋅ (*x*^2^ + *y*^2^ + *z*^2^), with *a* denoting a constant.

#### Trace segmentation

In the photon count trace segments, we distinguished single molecule emissions from background using a Hidden-Markov model^1^. We only considered trace sections for which the MINFLUX acquisition was running in its final iteration. We obtained a first estimate for the emission rates in the Hidden Markov model by calculating the mean emission after artificially splitting the median filtered photon counts at a manually chosen threshold in the range 1.1–3.6. We used a Poissonian distribution with the obtained mean emission values as initial emission probabilities in the Hidden-Markov model. Using the Matlab implementation of the Viterbi algorithm, we estimated the emission states from a three-state Hidden Markov model (1: on, 2: off, 3: blinking) with the sampling time *t*_s_=0.1 ms, *t*_off_=0.1 s, and *t*_on_=0.5 s being the estimated on and off-times of the molecule and *t*_blink,on_=1 ms and *t*_blink,off_=0.1 ms the estimated blinking on and off times. We applied the Viterbi algorithm twice, using the same transition probability matrix, but applying the improved emission rate distribution from the previous run. We merged successive emissions in states 1 or 3. We split events at a pre-defined photon number of *N*=2000 to obtain several localizations per molecule for an experimental assessment of the localization precision. We assigned a molecule identification number to each localization assuming that emissions with several iteration rounds (no activation applied in between) originated from the same molecule.

#### Position estimation

As previously reported, we used a maximum likelihood estimator implemented in a grid search optimization algorithm to retrieve the molecule positions from the photon count traces ^1^. For the 3D-MINFLUX scheme, we wrote the trivial extension of the *p*-functions to three dimensions using the presented TCP. Unlike before, we could not obtain the background counts directly from the measurement as the background depended on (i) the position in the scan and (ii) the position of the iterative TCP in the confocal volume, so that the background varied even for successive events of the same molecule. We circumvented this issue by estimating the signal-to-background ratio with the molecule position in the maximum likelihood estimator.

#### Event filtering

During the iterative MINFLUX measurement, we chose a fixed threshold value to decide on the presence of a molecule emission. This approach led to two types of false positives: (i) reaction to background, as we chose a low threshold value to avoid missing faint emission events; and (ii) reaction to thermally activated molecules outside the iterative MINFLUX region, which were still detected, despite the live estimators not reaching them.

We applied three filters in post-processing to select true emission events only. These are the central doughnut fraction *p*_0_, the estimated location of the molecule with respect to the center of the TCP *r*_rel_ and the photon number in the last iteration *N*. We show the distribution of the filtering variables together with the selected threshold values in figure Fig. S2–4 for all presented datasets.

Molecules far outside the TCP and background events are expected to produce equal mean counts in all *m* exposures, delivering 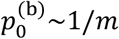, with 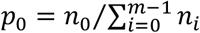 describing the probability to detect a photon in the central exposure. True emission events will yield *p*_0_ < 1/*m* when successfully approaching the molecule in the iterations. The overlap of the two distributions becomes stronger with decreasing SBR, so that a classification based on *p*_0_ is not sufficient in a cellular context. We achieved a better classification by using the distance of the estimated position with respect to the center of the TCP in the last iteration 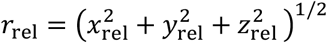, a measure that does not directly depend on the SBR. We note that further restricting both filter variables can improve the overall localization precision at the expense of discarding valid localizations. For measurements with a population of events with low photon number, we additionally applied a lower threshold on the photon number *N* in the last iteration to avoid a bias in the position estimation.

#### Multicolor classification

For each localization, we obtained photon numbers for both spectral channels in all iterations. We used the probability of detecting a photon in the blue-shifted spectral channel in the *i*-th iteration

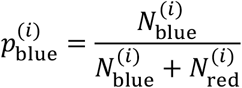

as a measure of the spectral properties of the dye. In the last iteration, we obtained mean values of 0.2, 0.4 and 0.55 for CF680, CF660C and Alexa Fluor 647 respectively. Photons from all MINFLUX iterations carry information about the spectral properties of the dye, but with different signal-to-background ratio values. To reduce the classification error by using all available information, we performed a principal component analysis based on 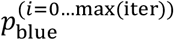 and manually chose a splitting threshold to classify the dye species based on the distribution of the first principal component.

#### Image rendering

In datasets with a low number of events per molecule, we replaced each localization with a Gaussian distribution, summing up pixel entries for overlapping Gaussians. We normalized the image, resulting in 0 ≤ *I*_*ij*_ ≤ 1 with *I*_*ij*_ being the value of the *ij*th pixel. For images displaying a micrometer sized image region (Fig. 2a, Fig. 5e), we chose a pixel size of 0.5 nm and a large width of the Gaussian kernel (*σ*=4 nm) for visibility. For images displaying single or few nuclear pores (Fig. 2a, f) we chose *σ*=2 nm and a pixel size of 0.2 nm. We used a non-linear color distribution to compensate for the unequal number of events per fluorescent molecule. For the multicolor data (Fig. 5e), we independently convolved the localization of each dye species with a Gaussian kernel and displayed the normalized images as RGB images with *α*=7 in the red channel and *α*=5 in the green channel using the components

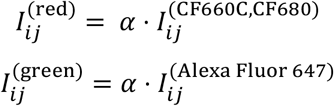

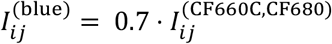.

For the DNA origami image (Fig. 5b), we displayed a simple 2D histogram of the localization data with 1 nm pixel size, with the RGB value for each pixel determined with *α*=0.5. Where we displayed localization data as scatter plots, the size of the scatter points/spheres was unrelated to the estimated localization precision. An exception is Fig. 2e, where we displayed scatter points together with an ellipse, with the principal axes lengths related to the localization precision.

#### Surface fitting

##### Nuclear pore complexes

To obtain a model of the nuclear envelope in the 3D measurements of nuclear pore complexes (Fig. 3f and Fig. 5e), we first determined the nuclear pore centers by clustering localization in the same nuclear pore. We employed the Matlab implementation *linkage* to generate an agglomerative hierarchical cluster tree using the median as a distance measure between clusters. We chose clusters using Matlab’s *cluster* function based on the manually determined number of clusters in the data. Based on the mean position of all clusters in the dataset we approximated the shape of the nuclear envelope based on thin-plate smoothing spline interpolation using a smoothing parameter of 0.999 (Matlab function *tpaps*).

##### PSD-95

Localizations were assigned to two domains using a *kmeans* clustering algorithm. To determine the average position of the volumes of high localization density within each domain, we employed the Matlab implementation of the density-based clustering algorithm *dbscan* (epsilon=8 nm, minPts=10). We determined a surface using the Matlab function *tpaps* with a smoothing parameter 0.996.

#### Performance metrics

##### Localization precision based on clustering

We assessed the localization precision in the acquired MINFLUX images based on a clustering of localizations according to their molecule identification number (see Trace segmentation). This included grouping localizations from single iteration cycles, obtained by splitting the photon numbers in time, as well as grouping localizations from subsequent iterations that were triggered by exceeding a pre-defined photon number in the last iteration. We filtered for molecules with more than four localizations. We calculated the standard deviation of single clusters and obtained the median standard deviation for all molecules (Fig. 2b, Fig. 3i). To achieve a more general assessment (Fig. 2c, Fig. 3h, Fig. 4c), we subtracted the mean cluster position from all localizations and displayed the obtained distances in 1D histograms for all three dimensions. We assumed a Gaussian distribution of the distances to determine their spread in a fit. We note that the calculation of the mean cluster position is biased for a low number of samples, which leads to a slight bias towards lower values in the localization precision that we obtain.

##### Fourier ring correlation

We used a Fourier ring correlation (FRC) analysis to assess the resolution in the image independently of any localization clustering. We randomly split the localizations in two groups and determined the FRC curve directly from the localization coordinates as previously described in ^5^. We chose a correlation threshold of 1/7 to determine the spatial frequency transition from signal-to-background (Nieuwenhuizen et al. 2013). To reduce noise in the determination of the cut-off frequency, we interpolated the data using the ‘nearest’ method of the Matlab function *interp1*.

### Sample preparation

#### Sample mounting and imaging buffers

To stabilize the sample during the MINFLUX acquisition, we treated all coverslips with gold nanorods (A12-25-980-CTAB-DIH-1-25, Nanopartz Inc., Loveland, CO, USA). We diluted the nanorods 1:3 in single molecule clean phosphate-buffered saline (PBS, P4417 Sigma-Aldrich, St. Louis, MO, USA), sonicated them for 5–10 min, and incubated the sample with the nanorod solution for 5–10 min at room temperature. To avoid floating nanorods, we washed the sample 3–4 times in PBS after removing the nanorod solution.

For MINFLUX imaging of samples labelled with Alexa Fluor 647, CF660C or CF680, we used a STORM blinking buffer containing 0.4 mg/mL glucose oxidase (Sigma, G2133), 64 µg/mL catalase (Sigma C100-50MG), 50 mM TRIS/HCl pH 8.0/8.5, 10 mM NaCl, 10–30 mM MEA (Cysteamine hydrochloride, Sigma M6500) and 10% (w/v) glucose. When measuring DNA origami samples, we added 10 mM MgCl_2_ to the blinking buffer to avoid dehybridization of DNA strands. We imaged U-2 OS cells endogenously tagged with mMaple in 50 mM Tris buffer (pH 8) in 95% D_2_O to reduce the short time blinking and increase the photon count of mMaple ^6^. We sealed the samples with picodent twinsil® speed 22 (picodent® Dental-Produktions-und Vertriebs-GmbH, Wipperfürth, Germany).

#### Fluorescent microsphere sample

We used samples with poly-L-lysin-immobilized fluorescent microspheres (FluoSheres®, 0.02 μm, dark red fluorescent; Thermo Fischer Scientific Inc., Waltham, MA, USA) to examine the setup performance on a daily basis, to measure the PSF of the optical setup and to calibrate the axial beam positioning with the VFL. The samples were prepared as previously described^1^.

#### DNA origami

We pre-annealed the unlabeled DNA origami template based on 10-fold excess of the staple strands (containing biotinylated strands for immobilization) compared to the scaffold strands (M13mp18, N4040S, New England Biolabs Inc., Ipswich, MA, USA) in folding buffer containing TAE 1x (1:5), MgCl_2_ 27 mM heating first to 85 °C for 3 min, cooling down at a speed of 0.5 °C/min until reaching 42 °C and storing at 4 °C. The sequences of the DNA staple strands are listed in the supplementary material (Tab. S2). After annealing, we purified the DNA origami templates by adding 15% PEG in TAE 1x containing 0.5 mM NaCl and 16 mM MgCl_2_. After mixing the solutions, we centrifuged for 30 min, keeping the sample at 4 °C. After removing excess, we repeated the procedure three times before storing the template in TAE 1x containing 10 mM MgCl_2_. Prior to experiments, we annealed the template with the complementary strands 5’-labelled with a fluorescent molecule. In a first step, we used a 25x excess of the labelled strands and kept the mixed solutions overnight at room temperature. After purifying as described above, we repeated annealing and purification with a 250x excess of labelled strands. On the day of the experiment, we prepared the samples with immobilized DNA origamis as previously described^1^.

#### U-2 OS Nup96-SNAP/mMaple

We handled the cells as previously described^7^. Briefly, cells were cultured in DMEM without phenol red (Cat#1180-02, ThermoFisher Scientific, Waltham, MA, USA) supplemented with 1x MEM NEAA (Cat#11140-035, ThermoFisher Scientific), ZellShield (Cat#13-0050, Minerva Biolabs, Berlin, Germany), 1x GlutaMAX (Cat#35050-038, ThermoFisher Scientific) and 10% [v/v] fetal bovine serum (Cat#10270-106, ThermoFisher Scientific) at 37 °C, 5% CO_2_ and 100% humidity. Seeded coverslips were pre-fixed (2.4% [w/v] formaldehyde [FA] in PBS) after two days of growth for 30 sec, permeabilized (0.4% [v/v] Triton X-100 in PBS) for 3 min and fixed (2.4% [w/v] FA in PBS) for 30 min. The sample was quenched by incubating it for 5 min in 100 mM NH_4_Cl and subsequently washed twice for 5 min in PBS. Samples of NUP96-mMaple were ready for imaging and mounted as described above. NUP96-SNAP samples were blocked for 30 min with Image-iT FX Signal Enhancer (ThermoFisher Scientific) and stained with staining solution (1 µM BG-AF647 [Cat#S9136S, New England Biolabs, Ipswich, MA, USA], 1 mM DTT in 0.5% [w/v] BSA) for 50 min. After washing 3 times in PBS for 5 min each to remove unbound dye, the samples were mounted as described above. For cells with an additional labelling for the center of the pore, we applied wheat germ agglutinin CF680 conjugate (29029, Biotium, Inc., Fremont, CA, USA) diluted to 0.02 µg/mL in PBS with an addition of 1% BSA in the final solution. We incubated the cells at room temperature for 5 min before washing three times in PBS. The cell lines are available via Cell Line Services (CLS, clsgmbh.de, Nup96-SNAP #300444, Nup96-mMaple #300461).

#### Hippocampal cultured neurons (PSD-95)

We prepared hippocampal neurons from transgenic PSD-95-HaloTag mice^8^ in accordance with the regulations of the German Animal Welfare Act and under the approval of the local veterinary service as previously described^9^. After 35 days in vitro we fixed the neurons in PFA (4% in PBS, pH 7.4) at room temperature for 15 min. After washing in PBS, we treated the neurons with Ammonium chloride (100 mM) for 10 min to reduce background due to autofluorescence. We permeabilized using 0.1% Triton X-100 in PBS for 10 min and subsequently stained with Alexa Fluor 647-HaloTag ligand (1 µM, synthesized in house) for 30 min at room temperature. After several short washing steps in PBS, we mounted the coverslips for imaging (see Sample mounting and imaging buffers).

## Data Availability

The data that support the findings of this study are available from the corresponding author upon reasonable request.

## Supplementary information

**Fig. S1.**
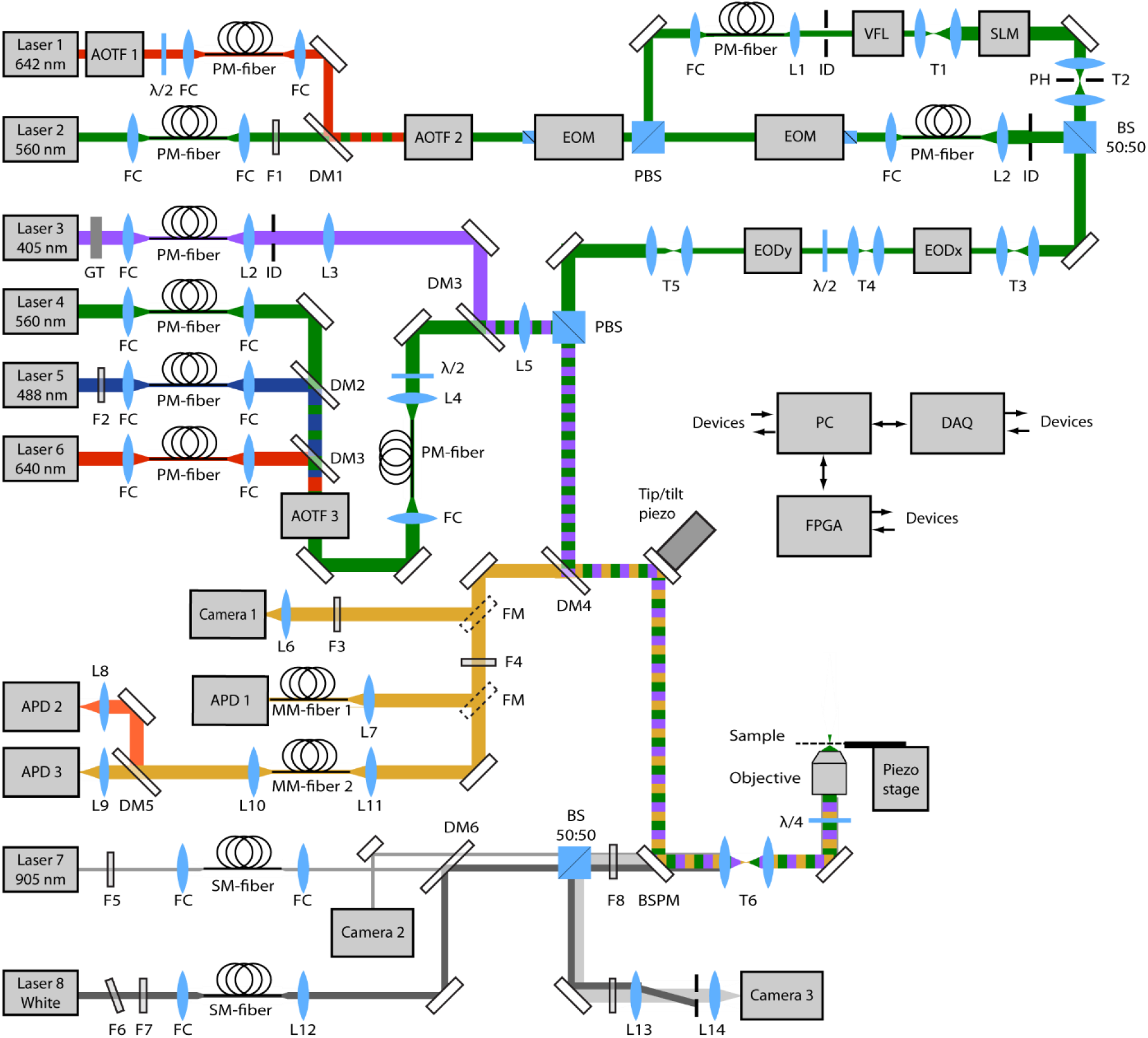
Schematic drawing of the optical setup. Lasers: Laser 1: VFL-P-1500-642 (MPB Communications Inc., Pointe-Claire, Quebec, Canada), Laser 2: Cobolt Jive™ 150-561 (Cobolt AB, Solna, Sweden), Laser 3: 405-50-COL-004 (Oxxius, Lannion, France), Laser 4: Cobolt Jive™ 25-561 (Cobolt AB, Solna, Sweden), Laser 5: LDH-D-C-485 (PicoQuant, Berlin, Germany), Laser 6: LDH-D-C-640 (PicoQuant, Berlin, Germany), Laser 7: LuxX® 905-150 (Omicron-Laserage Laserproodukte GmbH, Rodgau-Dudenhofen, Germany), Laser 8: Koheras SuperK Extreme (NKT Photonics, Birkerød, Denmark) Beam modulation: AOTF1: AOTFnC VIS-TN (AA Sa, Orsay, France), AOTF2: AOTFnC VIS-TN (AA Sa, Orsay, France), AOTF3: AOTFnC 400.650-TN (AA Sa, Orsay, France), EOM: LM 0202 P 5W + LIV 20 (Qioptiq Photonics GmbH & Co. KG, Göttingen, Germany), SLM: LCOS-SLM X13267-06 (Hamamatsu Photonics Deutschland GmbH, Herrsching am Ammersee, Germany), Scanning: EODx and EODy: M-311-A (Conoptics Inc., Danbury, CT, USA) + WMA-300 (Falco Systems BV, Amsterdam, The Netherlands), VFL: KLMS2D0700 −00 KTN varifocal lens module (NTT Advanced Technology Corporation, Omiya-cho Saiwai-ku, Kawasaki-shi, Japan) + AMPS-2B200-03 (Matsusada Precision Inc., Aojicho Kusatsu, Japan), Tip/tilt piezo: PSH-10/2 + EVD300 (both piezosystem jena GmbH, Jena, Germany), Piezo stage: P-733.3-DD + E725 (both Physik Instrumente (PI) GmbH & Co. KG, Karlsruhe, Germany), Polarization and beam transport: GT: Glan-Thompson polarizer (B. Halle Nachfl. GmbH, Berlin, Germany), PBS: polarizing beam splitter cube (B. Halle Nachfl. GmbH, Berlin, Germany), BS: beam splitter cube 50:50, FC: fiber collimator 60FC-* (Schäfter+Kirchhoff, Hamburg, Germany), λ/2: half wave plate (B. Halle Nachfl. GmbH, Berlin, Germany or Thorlabs Inc., Newton, NJ, USA), λ/4: achromatic quarter wave plate (Thorlabs Inc., Newton, NJ, USA), PM-fiber: polarization maintaining single mode fiber (Thorlabs Inc., Newton, NJ, USA or Schäfter+Kirchhoff, Hamburg, Germany), MM-fiber 1: multimode fiber M31L01 (Thorlabs Inc., Newton, NJ, USA), MM-fiber 2: multimode fiber M42L02 (Thorlabs Inc., Newton, NJ, USA), Lenses and mirrors Objective: HC PL APO 100x/1.40 Oil CS2 (Leica Microsystems GmbH, Wetzlar, Germany), L1-L14: achromatic lens with VIS or NIR AR coating (Thorlabs Inc., Newton, NJ, USA or Qioptiq Photonics GmbH & Co. KG, Göttingen, Germany), T1-T6: telescope, ID: iris diaphragm, FM: mirror on motorized flip mount, PH: pinhole, BSPM: back side polished mirror (Thorlabs Inc., Newton, NJ, USA), Dichroic mirrors and filters DM1: H 568 LPXR superflat (AHF Analysetechnik GmbH, Tübingen, Germany), DM2: Z500-RDC-XT (Chroma Technology Corp., Bellows Falls, VT, USA), DM3: Z620SPRDC (Chroma Technology Corp., Bellows Falls, VT, USA), DM4: ZT405/488/561/640rpc (AHF Analysetechnik GmbH, Tübingen, Germany), DM5: FF685-Di02 (Semrock Inc., Rochester, NY, USA), DM6: FF925-Di01 (Semrock Inc., Rochester, NY, USA), F1: ZET561/10x (Chroma Technology Corp., Bellows Falls, VT, USA), F2: 488/6 BrighLine HC (Semrock Inc., Rochester, NY, USA), F3: FF01-842/SP-25 (Semrock Inc., Rochester, NY, USA) and Quad-Band 446/523/600/677 HC (Semrock Inc., Rochester, NY, USA), F4: FF01-775/SP-25 (Semrock Inc., Rochester, NY, USA) and Quad-Notch 405/488/560/635 (Semrock Inc., Rochester, NY, USA) and ET700/75m (Chroma Technology Corp., Bellows Falls, VT, USA) or BLP02-561R-25 (Semrock Inc., Rochester, NY, USA), F5: FL905/10 (Dynasil, Littleton, MA, USA), F6: FELH0950 (Thorlabs Inc., Newton, NJ, USA), F7: FESH1000 (Thorlabs Inc., Newton, NJ, USA), F8: 66-230 long pass filter 950 (Edmund Optics®, Barrington, NJ; USA), Detectors APD 1: SPCM-AQR-13-FC (Excelitas Technologies, Waltham, MA, USA), APD 2,3: SPCM-AQRH-13-TR (Excelitas Technologies, Waltham, MA, USA), Camera 1: Ixon EMCCD DU897-BV, (Andor Technology Ltd., Belfast, UK), Camera 2: DMK 22BUC03 (The Imaging Source Europe GmbH, Bremen, Germany), Camera 3: DMK 23UP1300 (The Imaging Source Europe GmbH, Bremen, Germany), Computer PC: 3 personal computers running Windows 7 (Microsoft Corp., Redmond, WA, USA) and LabView 2016 (National Instruments, Austin, TX, USA), DAQ: NI PCIe-6353 + NI PCI-6259 (both National Instruments, Austin, TX, USA) + USB-3133 (Measurement Computing Corporation, Norton, MA, USA), FPGA: NI USB-7856R (National Instruments, Austin, TX, USA)

**Fig. S2.**
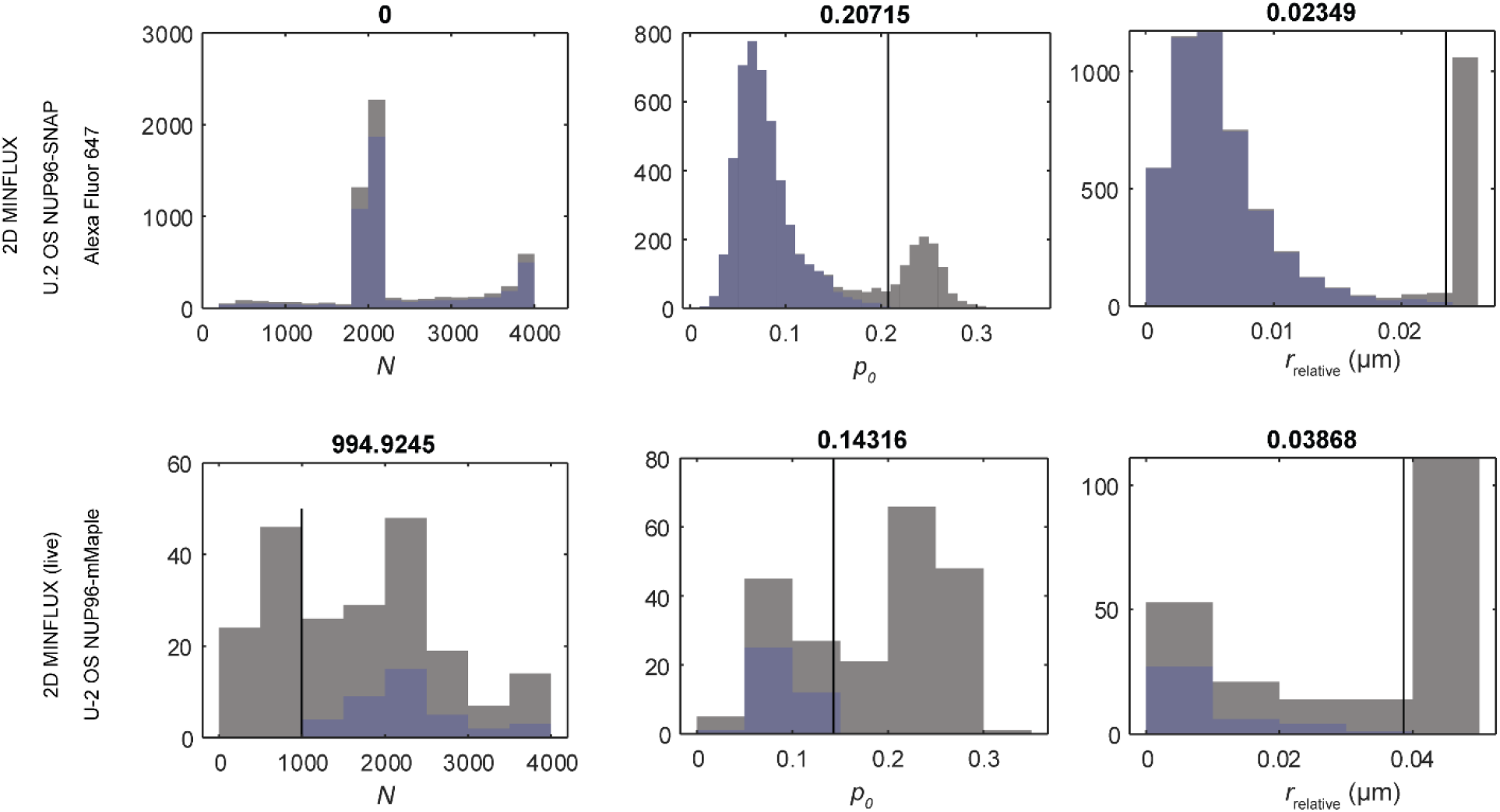
Filtering of 2D imaging data. The histogram of photon number *N*, the relative photon count number in the central exposure *p*_0_ and distance of the estimated position relative to the center of the last excitation beam pattern *r*_relative_ are displayed for each localization. Before filtering (gray) at a manually defined position (black line, number above), *p*_0_ and *r*_relative_ show two populations. The population that is assigned to background events is discarded, leading to a new filtered distribution (blue). Top row: data displayed in Fig. 2a. Bottom row: data displayed in Fig. 2f.

**Fig. S3.**
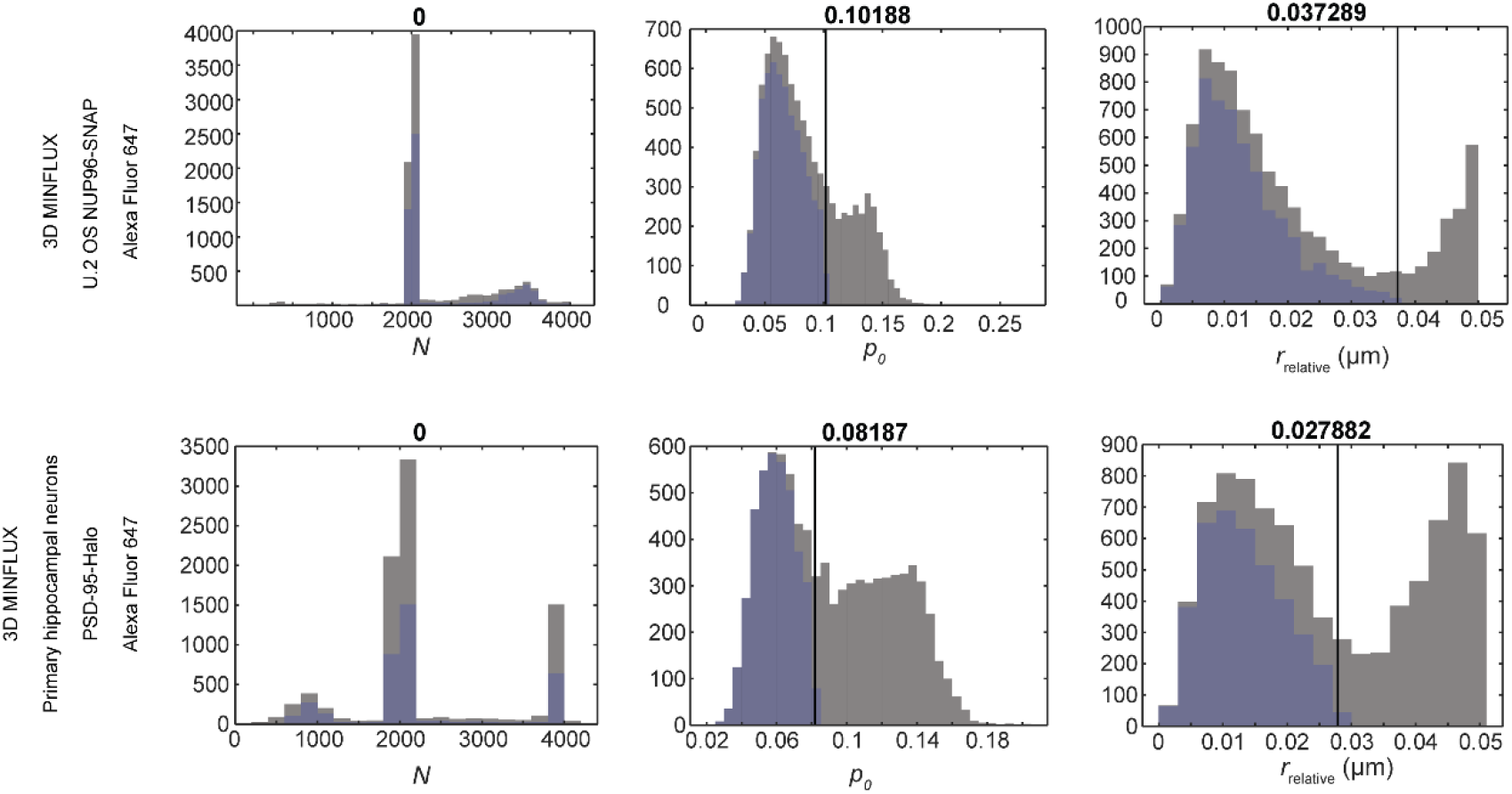
Filtering of 3D imaging data. The histogram of photon number *N*, the relative photon count number in the central exposure *p*_0_ and distance of the estimated position relative to the center of the last excitation beam pattern *r*_relative_ are displayed for each localization. Before filtering (gray) at a manually defined position (black line, number above), *p*_0_ and *r*_relative_ show two populations. The population that is assigned to background events is discarded, leading to a new filtered distribution (blue). Top row: data displayed in Fig. 3f. Bottom row: data displayed in Fig. 4a.

**Fig. S4.**
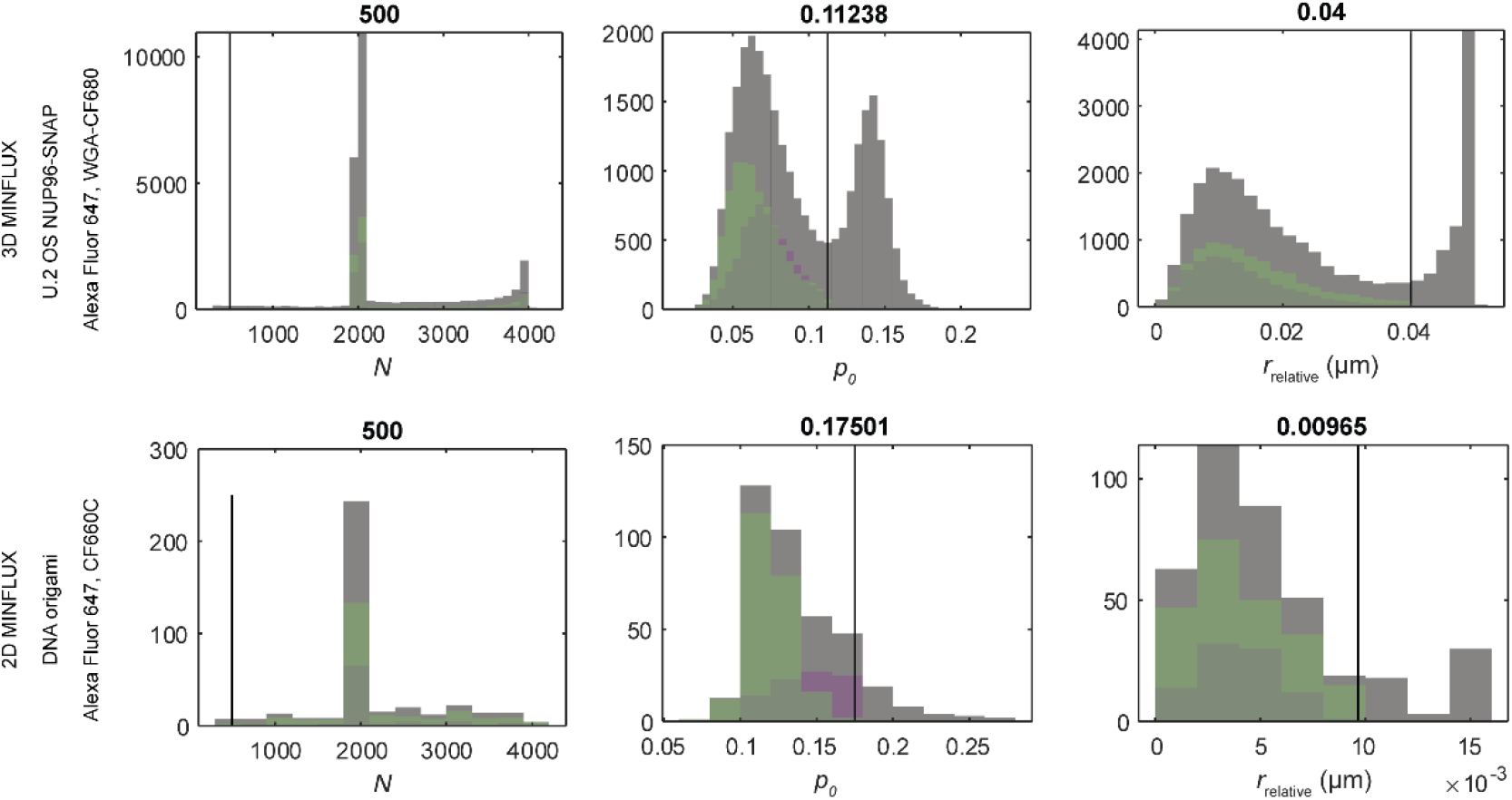
Filtering of multicolor imaging data. The histogram of photon number *N*, the relative photon count number in the central exposure *p*_0_ and distance of the estimated position relative to the center of the last excitation beam pattern *r*_relative_ are displayed for each localization. Before filtering (gray) at a manually defined position (black line, number above), *p*_0_ and *r*_relative_ show two populations. The population that is assigned to background events is discarded, leading to a new filtered distribution for each molecule species (green: Alexa Fluor 647, magenta: CF dye). Top row: data displayed in Fig. 5e. Bottom row: data displayed in Fig. 5c.

## Supplementary videos

**Video S1 | 3D MINFLUX nanoscopy of Nup96 in a mammalian cell.** U-2 OS cell expressing Nup96-SNAP labelled with Alexa Fluor 647 after fixation as displayed in Fig. 3. The color indicates the *z*-position of the localization.

**Video S2 | 3D MINFLUX nanoscopy of the synaptic protein PSD-95.** Primary hippocampal neurons from transgenic mice expressing PSD-95-Halo conjugated to Alexa Fluor 647 after fixation as displayed in Fig. 4. The color indicates the 3D localization density.

**Video S3 | 3D two-color MINFLUX nanoscopy of the nuclear pore complex in a mammalian cell.** U-2 OS cell expressing Nup96-SNAP labelled with Alexa Fluor 647 (green) and wheat germ agglutinin conjugated to CF680 (magenta) after fixation as displayed in Fig. 5. The colors indicate the molecular species assigned in the two-color classification.

---

**Tab. S1.**
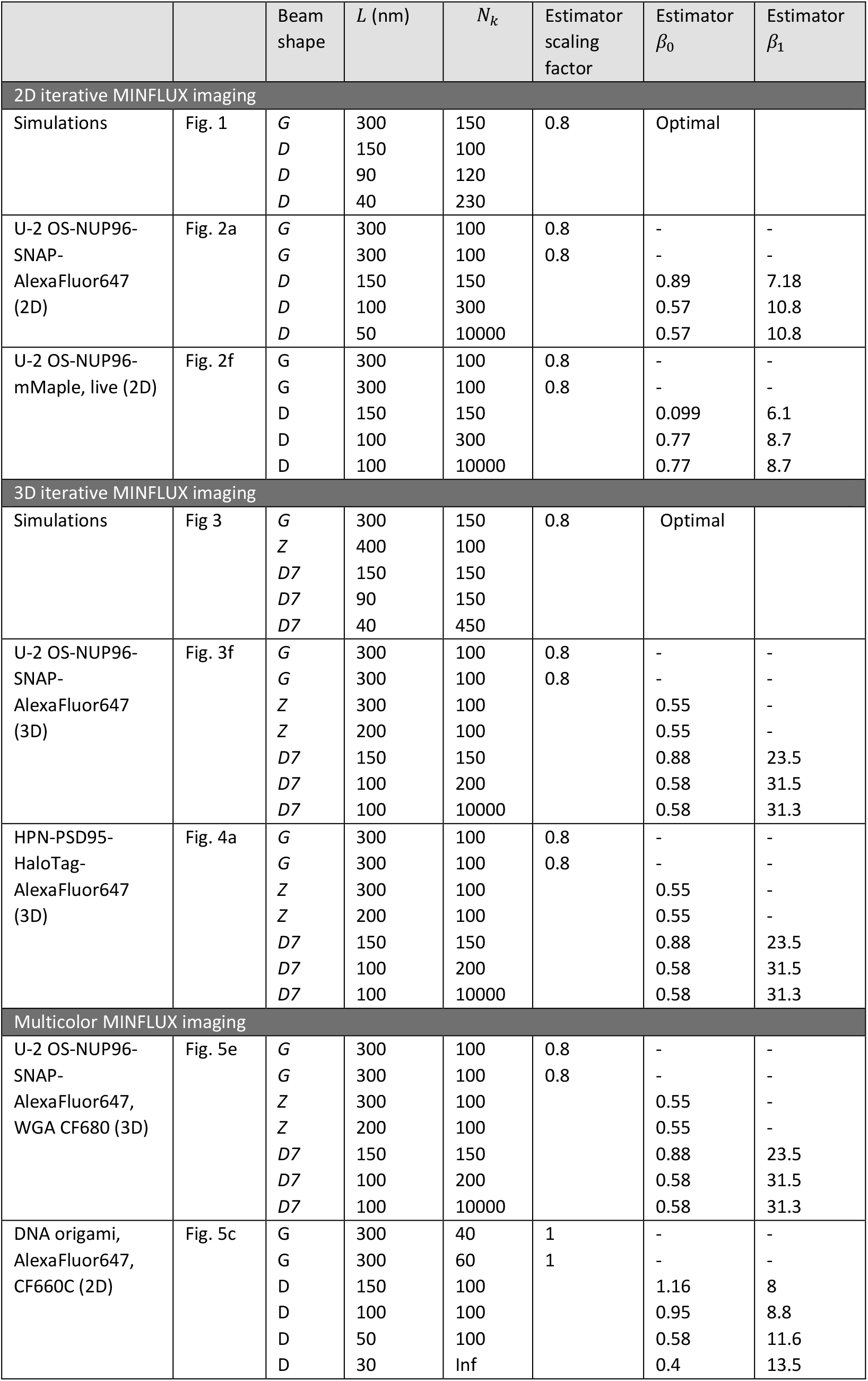
Iterative MINFLUX strategies

**Tab. S2.**
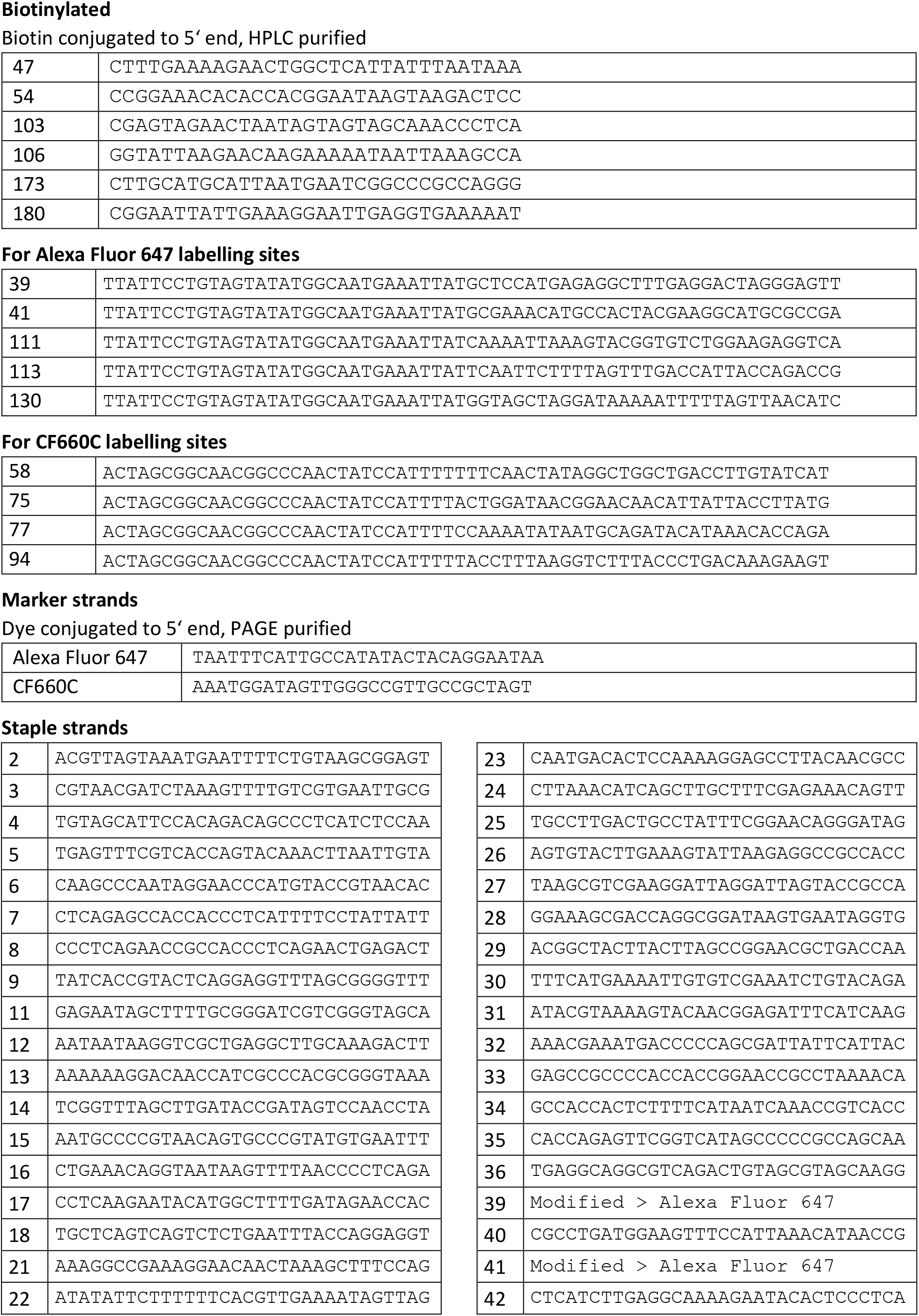
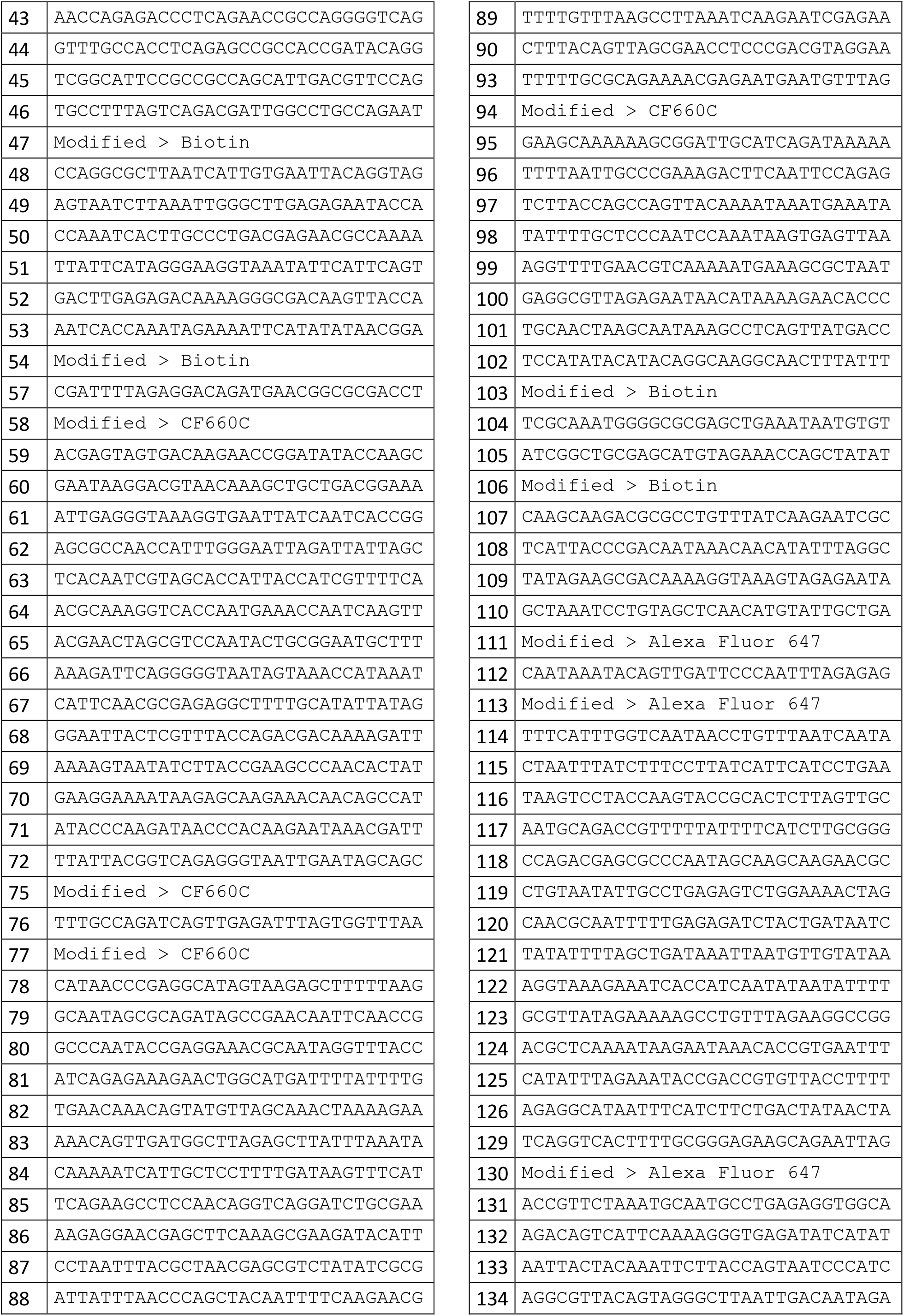
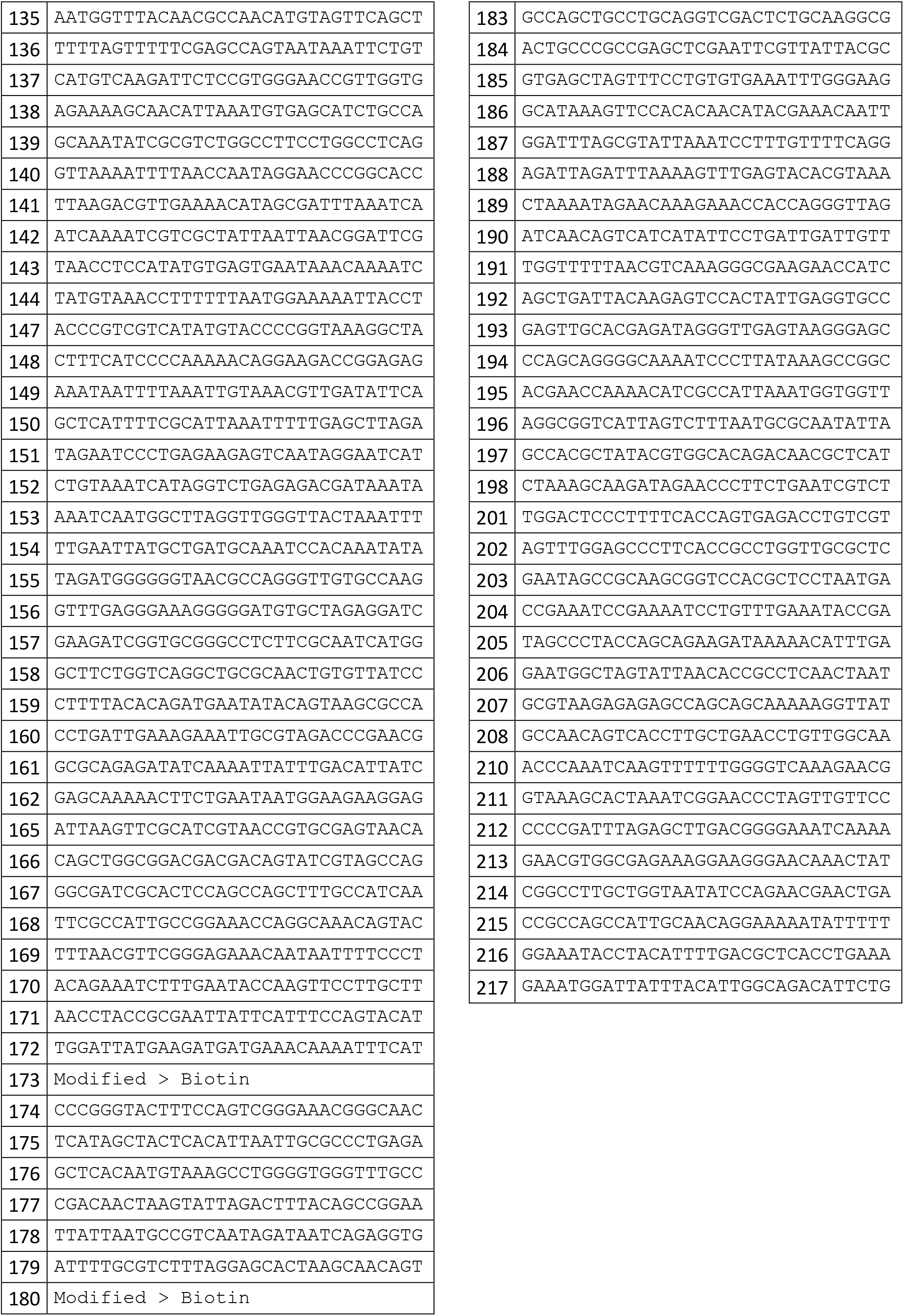
DNA origami strands.

## References

1. Hell, S.W. & Wichmann, J. Breaking the diffraction resolution limit by stimulated emission: stimulated-emission-depletion fluorescence microscopy. Opt Lett 19, 780–782 (1994).

2. Klar, T.A., Jakobs, S., Dyba, M., Egner, A. & Hell, S.W. Fluorescence microscopy with diffraction resolution barrier broken by stimulated emission. Proc. Natl. Acad. Sci. U.S.A. 97, 8206–8210 (2000).

3. Rust, M.J., Bates, M. & Zhuang, X. Sub-diffraction-limit imaging by stochastic optical reconstruction microscopy (STORM). Nat. Methods 3, 793–795 (2006).

4. Betzig, E. et al. Imaging Intracellular Fluorescent Proteins at Nanometer Resolution. Science 313, 1642–1645 (2006).

5. Balzarotti, F. et al. Nanometer resolution imaging and tracking of fluorescent molecules with minimal photon fluxes. Science 355, 606–612 (2017).

6. Hell, S.W. Far-Field Optical Nanoscopy. Science 316, 1153–1158 (2007).

7. Eilers, Y., Ta, H., Gwosch, K.C., Balzarotti, F. & Hell, S.W. MINFLUX monitors rapid molecular jumps with superior spatiotemporal resolution. Proc. Natl. Acad. Sci. U.S.A. 115, 6117–6122 (2018).

8. Backlund, M.P., Shechtman, Y. & Walsworth, R.L. Fundamental Precision Bounds for Three-Dimensional Optical Localization Microscopy with Poisson Statistics. Phys Rev Lett 121, 023904 (2018).

9. Thevathasan, J.V. et al. Nuclear pores as versatile reference standards for quantitative superresolution microscopy. bioRxiv, 582668 (2019).

10. von Appen, A. et al. In situ structural analysis of the human nuclear pore complex. Nature 526, 140 (2015).

11. Banterle, N., Bui, K.H., Lemke, E.A. & Beck, M. Fourier ring correlation as a resolution criterion for super-resolution microscopy. J Struct Biol 183, 363–367 (2013).

12. Nieuwenhuizen, R.P.J. et al. Measuring image resolution in optical nanoscopy. Nat. Methods 10, 557 (2013).

13. Hell, S.W., Reiner, G., Cremer, C. & Stelzer, E.H.K. Aberrations in confocal fluorescence microscopy induced by mismatches in refractive-index. J Microsc 169, 391–405 (1993).

14. Masch, J.-M. et al. Robust nanoscopy of a synaptic protein in living mice by organic-fluorophore labeling. Proc. Natl. Acad. Sci. U.S.A. 115, E8047–E8056 (2018).

15. MacGillavry, Harold, D., Song, Y., Raghavachari, S. & Blanpied, Thomas, A. Nanoscale Scaffolding Domains within the Postsynaptic Density Concentrate Synaptic AMPA Receptors. Neuron 78, 615–622 (2013).

16. Fukata, Y. et al. Local palmitoylation cycles define activity-regulated postsynaptic subdomains. The Journal of Cell Biology 202, 145–161 (2013).

17. MacGillavry, H.D. & Hoogenraad, C.C. The internal architecture of dendritic spines revealed by super-resolution imaging: What did we learn so far? Exp Cell Res 335, 180–186 (2015).

18. Zhang, Z., Kenny, S.J., Hauser, M., Li, W. & Xu, K. Ultrahigh-throughput single-molecule spectroscopy and spectrally resolved super-resolution microscopy. Nat. Methods 12, 935–938 (2015).

19. Li, B. & Kohler, J.J. Glycosylation of the Nuclear Pore. Traffic 15, 347–361 (2014).

20. Sharonov, A. & Hochstrasser, R.M. Wide-field subdiffraction imaging by accumulated binding of diffusing probes. Proc. Natl. Acad. Sci. U.S.A. 103, 18911–18916 (2006).

21. Jungmann, R. et al. Single-Molecule Kinetics and Super-Resolution Microscopy by Fluorescence Imaging of Transient Binding on DNA Origami. Nano Lett. 10, 4756–4761 (2010).

## References

1. Balzarotti, F. et al. Nanometer resolution imaging and tracking of fluorescent molecules with minimal photon fluxes. Science 355, 606–612 (2017).

2. Eilers, Y., Ta, H., Gwosch, K.C., Balzarotti, F. & Hell, S.W. MINFLUX monitors rapid molecular jumps with superior spatiotemporal resolution. Proc. Natl. Acad. Sci. U.S.A. 115, 6117–6122 (2018).

3. Gao, P., Prunsche, B., Zhou, L., Nienhaus, K. & Nienhaus, G.U. Background suppression in fluorescence nanoscopy with stimulated emission double depletion. Nat Photonics 11, 163 (2017).

4. Leutenegger, M., Rao, R., Leitgeb, R.A. & Lasser, T. Fast focus field calculations. Opt Express 14, 11277–11291 (2006).

5. Pham, T.-A., Soubies, E., Sage, D. & Unser, M. in ISBI 2019 - IEEE International Symposium on Biomedical Imaging (Venise, Italy;) 2019.

6. Ong, W.Q., Citron, Y.R., Schnitzbauer, J., Kamiyama, D. & Huang, B. Heavy water: a simple solution to increasing the brightness of fluorescent proteins in super-resolution imaging. Chemical communications (Cambridge, England) 51, 13451–13453 (2015).

7. Thevathasan, J.V. et al. Nuclear pores as versatile reference standards for quantitative superresolution microscopy. bioRxiv, 582668 (2019).

8. Masch, J.-M. et al. Robust nanoscopy of a synaptic protein in living mice by organic-fluorophore labeling. Proc. Natl. Acad. Sci. U.S.A. 115, E8047–E8056 (2018).

9. D’Este, E., Kamin, D., Göttfert, F., El-Hady, A. & Hell, S.W. STED Nanoscopy Reveals the Ubiquity of Subcortial Cytoskeleton Periodicity in Living Neurons. Cell Reports 10, 1246–1251 (2015).

